# X-TRUDE: A Process-Informed Framework for High-Fidelity Analysis of Hydrogel Extrusion

**DOI:** 10.64898/2026.02.16.706173

**Authors:** Farhad Sanaei, Pascal Bertsch, Juliette Lafosse, Sander C.G. Leeuwenburgh, Mani Diba

## Abstract

Reliable extrusion of viscoelastic hydrogels is crucial for technologies ranging from 3D (bio)printing to injectable therapeutics, yet current methods to characterize extrusion performance fail to mimic key processing conditions. Consequently, extrusion performance cannot be precisely predicted or controlled, particularly for emerging thermosensitive, shear-thinning, or heterogeneous hydrogels. Fundamentally, reliable extrusion is an emergent outcome of intrinsic materials properties coupled to extrinsic processing conditions.

Here, we introduce X-TRUDE, a process-informed characterization platform that recapitulates the spatiotemporal conditions of the extrusion process by integrating in situ pressure sensing, controlled thermal conditions, and process-relevant flow-path geometries. X-TRUDE reveals extrusion-specific phenomena inaccessible with conventional methods, including the transient evolution of apparent rheological response and time-local instabilities such as heterogeneity-induced variations in extrudate morphology. Across monolithic, thermosensitive, and granular hydrogel formulations, X-TRUDE establishes temporal pressure fluctuation patterns as quantitative metrics to unravel mechanisms underlying extrusion variability and correlate pressure profiles with extrudate morphology.

By linking intrinsic rheology with the physical extrusion environment, X-TRUDE provides a quantitative, mechanistic framework to benchmark extrudability across soft matter systems. This framework enables more reliable formulation development, reduces failure during process translation, and offers a generalizable tool for extrusion-based technologies in biofabrication, therapeutic delivery, and soft material manufacturing.

## 1. Introduction

Hydrogel extrusion plays a central role in various biomedical applications, ranging from advanced biofabrication techniques to minimally invasive treatments. Examples of these applications include extrusion-based 3D (bio)printing which has emerged as a leading technique for fabricating complex tissue-like constructs.^[1–4]^ Successful implementation of such extrusion-based processes critically depends on achieving consistent and reproducible extrusion which remains a major bottleneck. This limitation is often driven by flow instabilities during extrusion, which arise from the coupling between intrinsic materials properties and extrinsic processing conditions, producing emergent dynamic responses that evolve during extrusion.^[5–8]^ These challenges are further amplified for applications such as 3D (bio)printing and injectable hydrogel systems where the demands are increasing for novel formulations engineered to meet complex application-specific requirements.^[9]^ To overcome these barriers, a range of strategies has been explored through studying hydrogel rheology as well as thermal and geometric factors governing extrusion performance.

Rotational rheometry (RR) (**Figure 1a**) has long been employed as a widely used method for probing rheological behavior of hydrogels for extrusion-based applications.^[3,10–16]^ RR provides high-resolution viscometric data under idealized, wall-driven shear conditions.^[17]^ However, extrusion through a nozzle is governed by pressure-driven flow, resulting in spatially heterogeneous shear conditions (Figure 1b).^[18–21]^ Accordingly, by probing bulk material flow behavior, RR provides an incomplete basis for prediction of extrusion performance (Figure 1c), overlooking the transient variations in material response during extrusion (Figure 1d).^[10,18,22]^ Pressure-driven characterization approaches based on capillary rheology (CR) more closely resemble extrusion boundary conditions by probing flow through confined geometries and are well established in polymer processing.^[23,24,5]^ However, their adoption for hydrogel systems and biofabrication has remained limited.^[19,25–28]^ Force-based syringe measurements better approximate nozzle geometry yet merge intrinsic material response with system-dependent friction and pressure losses, yielding results that can vary substantially across hardware configurations.^[29–31]^ Microfluidic rheometry offers controlled, low-volume flow characterization, but omits the critical geometric transition from syringe to nozzle and thermal conditions happening in the actual process.^[32–34]^ In situ visualization methods are available, but remain focused on interrogating microstructural dynamics during flow and correlating these dynamics with bulk flow properties rather than providing a process-informed assessment of extrusion performance.^[35–38]^ Finally, retrospective characterization based on post-process quality checks, such as visual inspection of extrudates and/or intended constructs, provide only semi-quantitative information on extrusion performance.^[39–41]^

**Figure 1.**
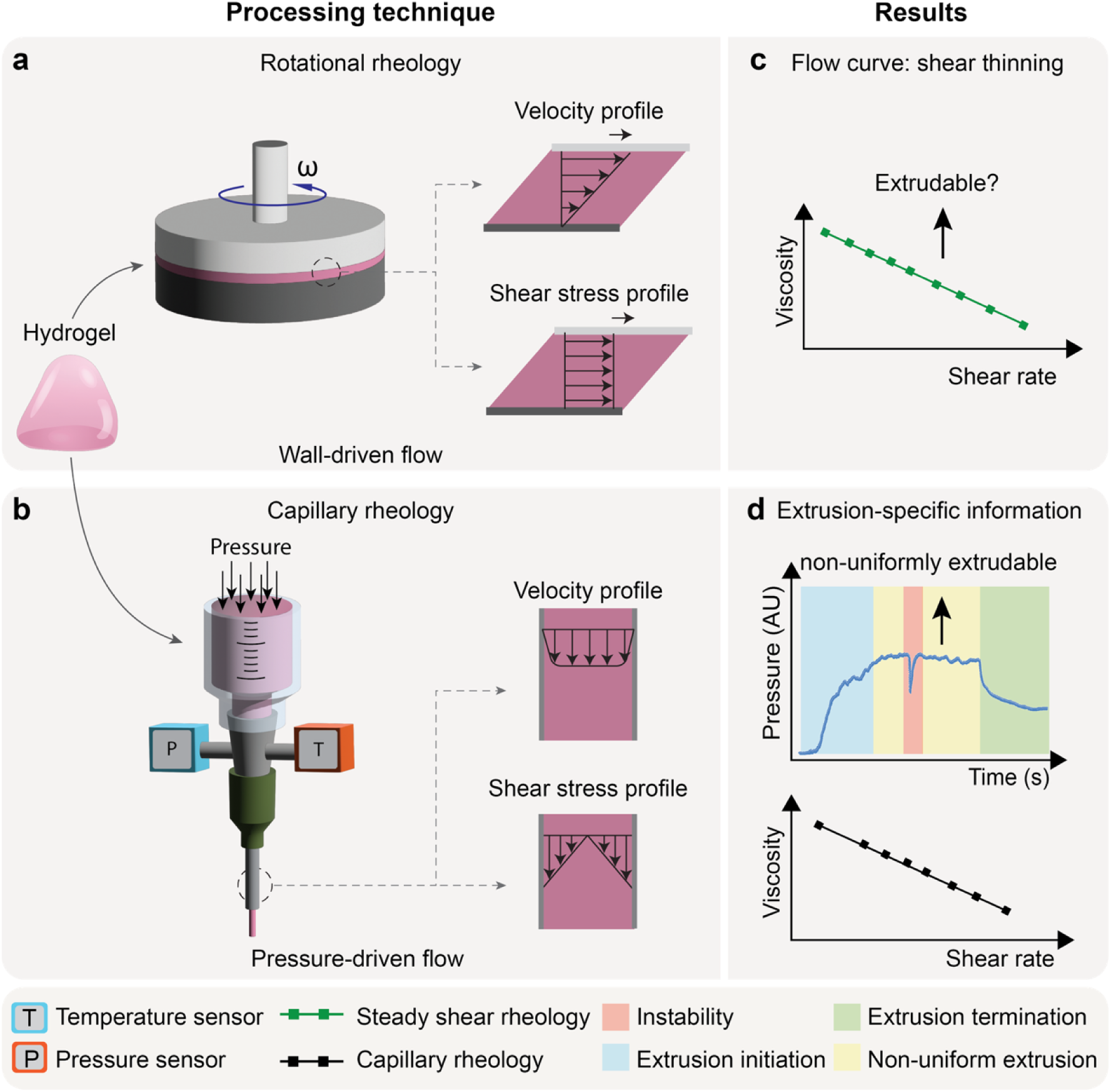
Schematic comparison of hydrogel extrusion characterization by means of rotational rheology (RR) using a conventional rheometer versus capillary rheology (CR) using the X-TRUDE system. (a) Steady shear characterization of a hydrogel using RR, with velocity and shear stress profiles across the sample gap generated by wall-driven flow. (b) X-TRUDE assessment of the same material, with velocity and shear stress profiles generated by pressure-driven flow, captured through simultaneous in situ recording of pressure and temperature (c) Representative flow curve (i.e., viscosity versus shear rate) obtained by RR. (d) Representative pressure-time response and flow curves obtained by CR using X-TRUDE; highlighted regions denote emergent extrusion-specific dynamic responses of the hydrogel.

Collectively, these approaches represent important advances towards characterizing hydrogel extrusion by offering valuable insights into bulk rheology, capillary flow, or microstructural dynamics. These advances, however, capture only isolated aspects of extrusion process and thus fall short of establishing a predictive framework under application-relevant condition. This persistent gap continues to constrain the rational design and translation of hydrogel formulations for extrusion-driven technologies, including extrusion-based 3D (bio)printing. It therefore remains a critical challenge to develop a characterization platform that directly couples intrinsic hydrogel properties with extrusion-relevant flow behavior under realistic processing environments.

To address this critical gap, here we introduce X-TRUDE (Extrusion Rheology for Understanding and Defining Extrudability). Unlike conventional techniques, X-TRUDE probes intrinsic materials properties within extrinsic processing conditions and captures their interaction under extrusion-relevant environments. By faithfully recapitulating flow dynamics, thermal environment, and geometric constraints of extrusion systems, X-TRUDE enables simultaneous, real-time quantification of materials properties—including viscosity and shear-thinning behavior—as well as emergent dynamic responses such as temporal flow stability and extrudate uniformity. Through this integrated approach, X-TRUDE bridges the gap between standard rheological analysis and actual extrusion performance, establishing a mechanistic foundation for the rational design and translation of hydrogel formulations across extrusion-driven technologies.

## 2. Results and Discussion

The X-TRUDE system was developed as a temperature-controlled capillary rheology platform that enables quantitative, in situ analysis of hydrogel extrusion behavior under process-relevant conditions. An overview of the system and its principal analytical features is presented in **Figure 2**, including real-time in situ pressure/temperature probing, geometrical constraints, and quantification of rheological and transient information. Additional details on system design, and measurement principles are provided in **Figure S1-S13**. Each of these features is discussed in detail in the subsequent sections, where their relevance to extrusion behavior and printability assessment is systematically evaluated. The X-TRUDE system consists of a temperature-controlled extrusion chamber (Figure 2a) that enables simultaneous extrusion of two syringes, while independently monitoring extrusion pressure and temperature in situ. Use of two syringes with different capillary length allow for downstream correction for calculation of hydrogel viscosity (SI section 2.3). The strategic integration of sensors directly within the flow path enables real-time acquisition of dynamic flow behavior, which is not feasible with RR. By accounting for multiple extrusion-related parameters, such as extrusion temperature, pressure, and geometrical features (Figure 2b), the X-TRUDE captures intrinsic rheological characteristics under extrusion (Figure 2c) as well as process-dependent phenomena. These transient phenomena encompass multiple layers of spatiotemporal material responses, including thermal gradients commonly encountered in 3D (bio)printing (Figure 2d) and phase separation of hydrogels (i.e., filter-pressing) during extrusion (Figure 2e). These capabilities allow for accurate prediction of material behavior under application-relevant extrusion conditions. Moreover, X-TRUDE facilitates advanced analyses, including inferring extrudate morphology solely from in situ pressure data without requiring visual monitoring of the extrudate (Figure 2f), highlighting its ability to extend beyond conventional rheological measurements.

**Figure 2.**
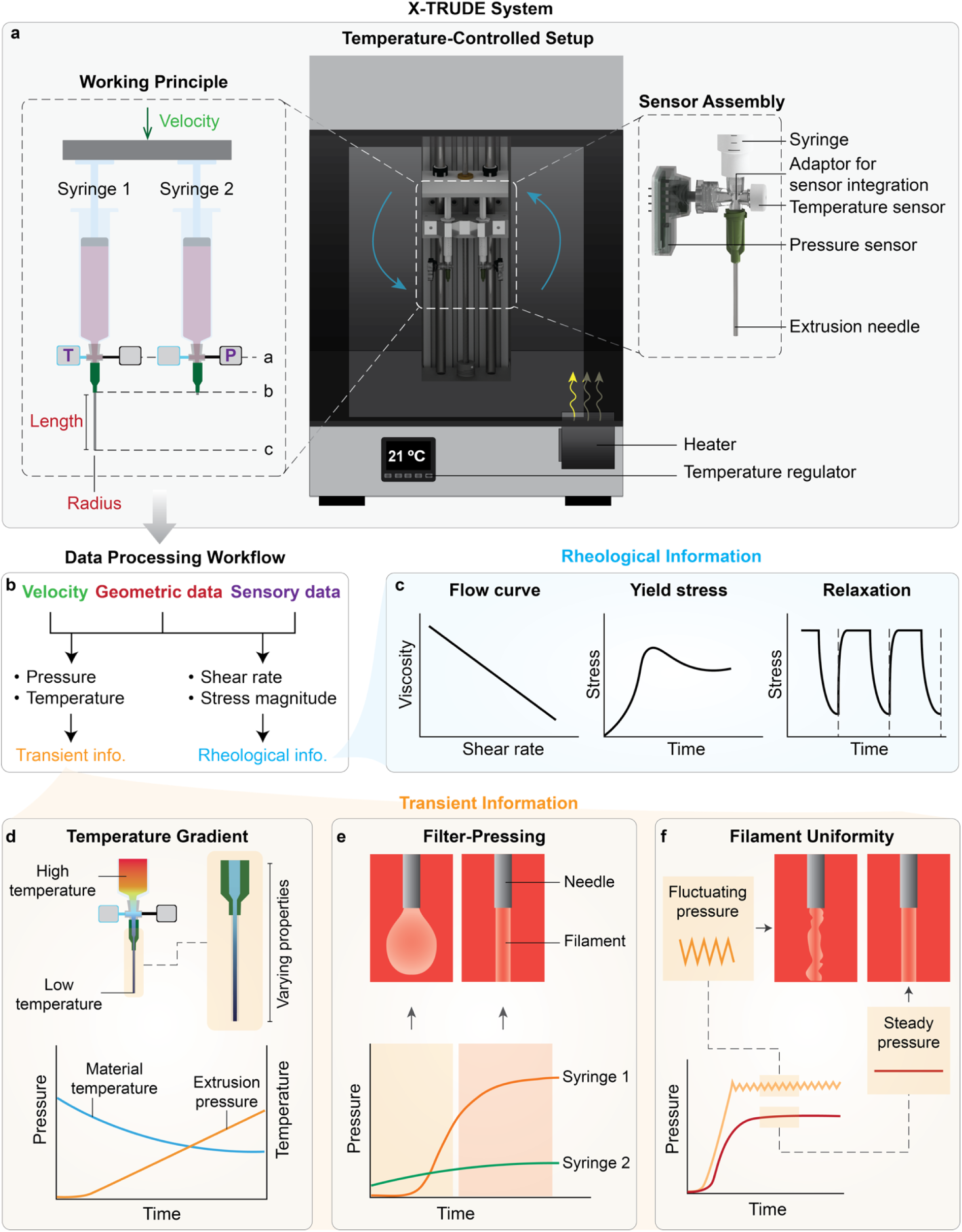
Schematic overview of the X-TRUDE system, its working principle, associated data-processing workflow and key features. (a) General layout of the X-TRUDE setup, including a 3D CAD view of the temperature-controlled chamber, thermal control unit, and the arrangement of syringes and sensors on the extrusion unit, alongside a schematic illustrating the working principle of the system. Yellow and blue arrows in the CAD view indicate temperature input from the heater and air recirculation within the chamber, respectively. The working-principle schematic shows two syringes connected to capillaries of defined length and inner radius, equipped with pressure sensors (black) and temperature sensors (blue) positioned at the level denoted by dashed line “a”. T indicates the in situ temperature of the material inside the capillary. Both syringes are actuated by a unified plunger at a constant linear velocity. Dashed line “b” denotes pressures at the inlet of both capillaries, and “c” denotes the pressure at the tip of the capillary on syringe 1. (b) Data-processing workflow outlining how input extrusion velocity, pressure and temperature sensory data, and geometrical data (i.e., length and inner radius of capillary) are used to extract rheological and transient information. (c) Illustration of representative rheological outputs of the X-TRUDE system, including flow curves (viscosity vs. shear rate), yield-stress identification, and stress-relaxation profiles. (d-f) Representative illustration of key quantitative transient outputs measured by X-TRUDE system. (d) characterization of spatiotemporal temperature gradients within the capillary, with corresponding time-dependent thermorheological changes reflected in extrusion pressure response of material extruded through syringe 1. e) Filter-pressing schematic showing the temporal pressure development of syringe 1 and syringe 2, alongside their corresponding extrudate morphology. This phenomenon divides the pressure response into an initial phase, where the syringe 1 extrusion pressure is lower than that of syringe 2, and a later phase, where syringe 1 extrusion pressure is higher than that of syringe 2. f) Filament uniformity assessment based solely on in situ pressure measurement, demonstrating a non-uniform filament associated with pronounced pressure fluctuations (left) compared to a uniform filament with minimal pressure fluctuations (right), alongside their representative filament schematics.

To quantitatively assess the predictive performance and added value of X-TRUDE relative to conventional RR, we selected three hydrogel formulations representing distinct classes of materials commonly employed in extrusion-based applications. Pluronic F-127 was used as a model synthetic shear-thinning hydrogel,^[40,42]^ monolithic gelatin methacryloyl (mGelMA) as a homogeneous natural polymer hydrogel,^[3,43]^ and granular GelMA (gGelMA) as a representative jammed microgel formulation.^[43,44]^ While mGelMA is capable of undergoing covalent photo-crosslinking, the formulations used in this study were not crosslinked in order to reflect their commonly employed state prior to extrusion. Nevertheless, the mGelMA precursor solution exhibited physical gelation at room temperature or below due to the intrinsic thermally-induced physical network assembly of gelatin.^[3]^ Moreover, although the hydrogel fragments used as building blocks for gGelMA were covalently photo-crosslinked during their initial fabrication, the resulting granular formulation relied primarily on physical jamming and non-covalent interactions between the hydrogel fragments.

We first focused on mGelMA to establish the methodological correspondence and divergence between X-TRUDE and RR under controlled, steady-state conditions. We then extended the X-TRUDE analysis to all three material systems to resolve extrusion-specific, emergent dynamic responses such as filter-pressing and extrusion-path clogging. Finally, temperature-dependent and morphology-predictive studies were conducted using mGelMA as a representative thermosensitive hydrogel to elucidate the broader process parameters influencing extrusion fidelity.

### 2.1 Extrusion performance of hydrogels emerges from material-process coupling

Shear-thinning, characterized by a reduction in viscosity with increasing shear rate, is widely considered essential for facilitating the extrusion of hydrogels through narrow nozzles under cell-compatible stress conditions.^[45,46]^ In practice, materials exhibiting pronounced shear-thinning are often classified as injectable or printable based solely on this behavior, with the underlying assumption that reduced viscosity under shear directly translates to smooth extrusion. This assumption is common in biofabrication research, where steady-shear flow sweeps obtained by RR are frequently interpreted as indicators of extrudability.^[10,12,15]^

To investigate how reliably shear-thinning behavior as examined by RR predicts extrudability, we used mGelMA as a representative homogeneous hydrogel system commonly used as a bio-ink in bioprinting and as a biomaterial in minimally invasive treatments in preclinical phase.^[47–49]^ Two widely used formulations—5 wt% (mGelMA-5) and 10 wt% (mGelMA-10)—were first selected to evaluate how variations in polymer content influence rheological and extrusion behavior. To compare RR with X-TRUDE, all formulations were prepared using GelMA from the same supplier batch to ensure consistent physicochemical characteristics. This controlled design ensured that any observed differences reflect methodological rather than compositional effects. Steady-shear flow sweep tests performed with RR at 21 °C confirmed pronounced shear-thinning behavior in both formulations. For both formulations, viscosity decreased by approximately four orders of magnitude over a shear-rate range of 0.1–100 s⁻¹ (**Figure 3a**). Compared with RR, X-TRUDE revealed significantly different flow behavior for the same materials and, importantly, distinguished between the extrusion responses of mGelMA-5 and mGelMA-10 under pressure-driven conditions (Figure 3b). During the extrusion process, the pressure values for mGelMA-5 increased from 0 mbar and stabilized at 1327 ± 181 mbar, indicating a balance between applied pressure and resistance of mGelMA-5 to flow. In contrast, pressure values for mGelMA-10 rose rapidly without reaching a steady plateau before the test was manually stopped at ∼8 bar to prevent exceeding safe operational limits of the syringe (8.6 bar, ISO 7886-1:2017^[50]^ per manufacturer specifications). These results confirm that mGelMA-10 was not extrudable under the tested conditions (21 °C, 18G nozzle, 10 mm s⁻¹, ∼95 s⁻¹). Based on microscopic observations of the nozzle tip, when extruded through an 18G capillary (inner diameter = 0.84 mm), mGelMA-5 produced inconsistent, non-uniform flow, while mGelMA-10 failed to extrude entirely (Figure 3c). These findings emphasize that shear-thinning in RR alone does not reliably predict extrudability. The discrepancies observed between RR and X-TRUDE measurements arise from the combined influence of both material and process-dependent factors that govern flow stability.

**Figure 3.**
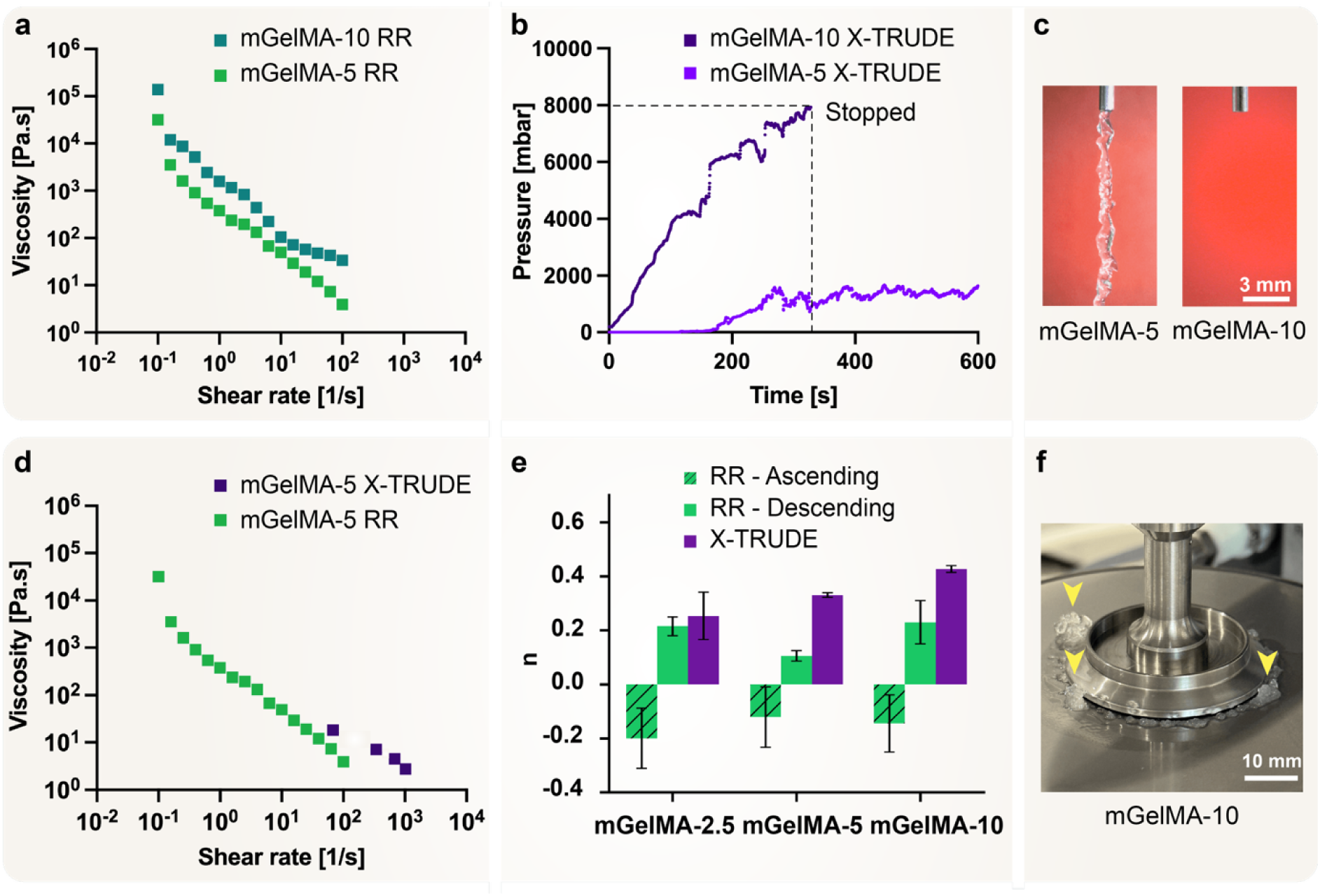
Rheological behavior of mGelMA formulations characterized using rotational rheology (RR) and the X-TRUDE system. (a–c) Flow behavior assessed (a) using RR shown as flow curves, (b) using X-TRUDE shown as pressure-time plots (mGelMA-10 stopped at syringe pressure tolerance), and (c) representative images of the nozzle tip/extruded filament. X-TRUDE measurements (b-c) were performed using an 18G nozzle at 10 mm s⁻¹. (d–f) Flow behavior inconsistencies shown as (d) comparative RR and X-TRUDE flow curves, (e) shear-thinning index *n* from RR (ascending: 0.1–100 s⁻¹; descending: 100–0.1 s⁻¹) and X-TRUDE (∼1–100 s⁻¹ for mGelMA-10; ∼1–130 s⁻¹ for mGelMA-5), and (f) material expulsion from geometry-plate gap (indicated with yellow arrows) during flow sweep characterization. Data in panel (e) are presented as mean ± standard deviation (n = 2 for RR, n = 3 for X-TRUDE). All experiments were carried out at room temperature (21 °C).

Previous studies suggest that these discrepancies arise from the interplay between yield stress, time-dependent viscoelastic behavior, and temperature rather than shear-thinning alone.^[41,46,51]^ Compositions exhibiting low yield stresses were shown to lead to droplet-like, or unstable strands formation, and compositions with too high yield stress were not extrudable.^[51]^ Factors such as material composition and viscosity were crucial as well, with extrudability emerging only when these parameters jointly fall within a certain viscosity range.^[52]^

Beyond these mechanistic considerations, our results further illustrate how differences in the underlying flow mechanisms between RR and X-TRUDE influence the accuracy of viscosity measurements. For example, the flow curve of mGelMA-5 determined by X-TRUDE showed viscosity values approximately threefold higher than those measured by RR across comparable shear-rate ranges (Figure 3d). Obtaining flow curves for mGelMA-10 was technically unattainable using the X-TRUDE system, as the material did not exit the nozzle at pressure ranges below syringe tolerance. This mismatch is in line with previously reported findings for other material formulations, where formulations with high polymer content were not extrudable, despite showing shear-thinning behavior when characterized using RR.^[19]^ RR is characterized by wall-driven flow, in which the material is placed between two surfaces with one or both rotating to impose shear deformation. Conversely, extrusion through capillary involves pressure-driven extrusion, where the material flows through a narrow capillary channel, experiencing a combination of shear and elongational stresses.^[53]^

Due to these fundamental process differences, the flow in RR is predominantly shear-driven with no elongational components. Consequently, the viscosity measured via RR primarily reflects the response of materials to pure shear conditions.^[17]^ In contrast, X-TRUDE measurements inherently combine shear with substantial elongational flow, particularly pronounced at the capillary entrance and exit regions.^[18,20]^ Such mixed flow conditions in X-TRUDE lead to viscosity profiles that differ from those obtained by RR, reflecting a more realistic representation of the material’s resistance under extrusion-relevant flow conditions. Specifically, the elongational stresses present in X-TRUDE contribute an additional resistance to flow via an extensional component, a phenomenon particularly pronounced in viscoelastic materials.^[18,21]^

These mechanistic differences also determine the range and magnitude of shear rates that can be realistically achieved by each method. X-TRUDE closely replicates the shear profiles and magnitudes typical of hydrogel extrusion processes. For instance, analysis of literature reports on extrusion-based 3D (bio)printing of hydrogels indicates a median shear rate of ∼121 s⁻¹ (interquartile range: 43–267 s⁻¹; **Figure S14**). X-TRUDE encompasses this typical shear rate range showing a lower operational limit of ∼1 s⁻¹ defined by the instrumental pressure-transducer resolution at low flow rates and a higher limit of well up to ∼1000 s⁻¹ depending on material formulation and viscosity. In contrast, while RR provides reliable viscosity data at low shear rates (0.01–100 s⁻¹), measurements above ∼100 s⁻¹ become unreliable, as fluid inertia, detachment or fracture of the sample at the geometry edge, and sample ejection often arise,^[22]^ leading to erroneous viscosity values.

A further discrepancy was observed within RR-derived flow curves when calculating power law flow behavior index (n), which is a commonly used index to quantify shear-thinning behavior.^[54,55]^ Specifically, n values were negative (n_mGelMA-2.5_ = -0.21 ± 0.03, n_mGelMA-5_ = -0.20 ± 0.01, n_mGelMA-10_ = -0.14 ± 0.11) when applying flow sweeps with ascending shear rate (0.1 to 100 s^-1^), indicating an artifact-driven response inconsistent with physical flow behavior (Figure 3e). In contrast, flow sweep with descending shear rates (100 to 0.1 s⁻¹) yielded positive n values (n_mGelMA-2.5_ = 0.22 ± 0.04, n_mGelMA-5_ = 0.11 ± 0.02, n_mGelMA-10_ = 0.23 ± 0.08) which were consistent with literature reports.^[10]^ While negative flow indices under ascending shear have been reported for some materials,^[54]^ such protocol-dependent outcomes emphasize a need for careful evaluation of RR-derived results. Conversely, X-TRUDE measurements revealed positive n values for both mGelMA compositions. However, increasing n values were observed with increasing polymer concentration (n_mGelMA-2.5_ = 0.25 ± 0.09, n_mGelMA-5_ = 0.33 ± 0.01, n_mGelMA-10_ = 0.43 ± 0.01) in contrast to trends observed in RR and in literature reports.^[10,54,55]^ The occurrence of non-physical (negative) n-values in ascending shear sweeps suggested potential measurement artifacts arising from sample instability. To further investigate this possibility, we examined RR flow curves for evidence of material expulsion or deformation-induced artifacts. This analysis revealed a sharp initial drop in viscosity values at the onset of shear in both ascending and descending flow sweeps (Figure 3a, **Figure S15.**). This phenomenon was visually correlated with the expulsion of material from the geometry-plate gap (Figure 3f, **Figure S16**), which can significantly compromise the reliability of RR for flow behavior analysis of hydrogels.

To gain deeper insight into the material expulsion during RR measurements, we next analyzed axial force values recorded during shear sweep measurements, as a key complementary parameter for interpreting RR results. These evaluations revealed a notably elevated levels of normal force under ascending shear (N_mGelMA-B2-10_ = 41.3 N, N_mGelMA-B2-5_ = 12.6 N, N_mGelMA-B2-2.5_ = 0.5 N, n = 1). In contrast, descending shear resulted in substantially lower normal force values (N_mGelMA-B2-10_ = 7.2 N, N_mGelMA-B2-5_ = 5.4 N, N_mGelMA-B2-2.5_ = 0.08 N, n = 1) (**Figure S17**).

The occurrence of sample expulsion even at shear rates as low as 0.16 s⁻¹ suggests that the phenomenon arises from brittle failure, where applied stress induces fracture rather than uniform flow after yielding. Consequently, although geometries such as cup-and-bob could physically contain the sample, they would not eliminate this intrinsic failure mechanism, which fundamentally limits the applicability of RR for analyzing the flow behavior of viscoelastic hydrogels.^[45]^ Nevertheless, further investigations are required to elucidate the detailed mechanisms underlying this failure behavior, which are beyond the scope of the present study. While the preceding analysis highlights measurement artifacts intrinsic to RR, variability in the material itself can also influence extrusion behavior. In particular, natural polymers such as gelatin are known to commonly exhibit batch-to-batch variability arising from intrinsic structural, and compositional heterogeneity inherent to biologically derived macromolecules.^[56]^ Having established the methodological differences between RR and X-TRUDE using GelMA from a single batch, we next examined whether these techniques could detect subtle material variations that can naturally occur between different GelMA batches. To evaluate whether RR and X-TRUDE are sensitive to capture such potential variations, a second batch of monolithic GelMA (mGelMA-B2; produced under the same formulation and quality specifications as the first batch) was analyzed. RR results were consistent with the previous batch, with each composition in the new batch (mGelMA-2.5-B2, mGelMA-5-B2, and mGelMA-10-B2) showing similar “shear-thinning” behavior to the corresponding composition in previous batch (Figure S15). However, extrusion behavior measured by X-TRUDE diverged significantly between the two batches. Notably, mGelMA-10-B2 extruded at 21 °C under 4 bar pressure (**Figure S18**), whereas the same formulation from the first batch failed to extrude even at 8 bar. These results indicate that minor batch-to-batch variations can exert a deterministic impact on the extrusion behavior of hydrogels, yet can remain undetectable in conventional RR, while X-TRUDE can capture such differences more faithfully by recapitulating realistic processing conditions.

Taken together, these findings emphasize the limited translational relevance of RR-based flow behavior measurements to application-relevant extrusion processes. The divergence between RR and X-TRUDE results underscores the process-dependency of extrusion and highlights importance of using process-informed methods when evaluating hydrogel materials such as GelMA in extrusion-based workflows.

### 2.2. X-TRUDE effectively identifies extrusion-specific emergent dynamic responses

Having established the process-dependent mechanistic differences between RR and X-TRUDE using mGelMA, we next examined whether X-TRUDE could resolve time-dependent extrusion phenomena that arise during continuous flow. Conventional rheological methods fail to capture key transient features that arise during extrusion such as pressure build-up, filament instabilities, or hydrogel–geometry interactions, which are critical determinants of reliable extrudability, particularly in 3D (bio)printing applications.^[3,11,15,42,46,54]^ To address these limitations, we next employed X-TRUDE system for real-time monitoring of pressure evolution and extrudate morphology under process-relevant conditions (**Figure 4**). To this end, we extended our analysis to three representative categories of hydrogels, i.e., 25 wt% Pluronic F-127 (Pluronic-25), 2.5 wt% mGelMA (mGelMA-2.5), and granular GelMA (gGelMA, fragmented and jammed form of 5 wt% crosslinked GelMA).

**Figure 4.**
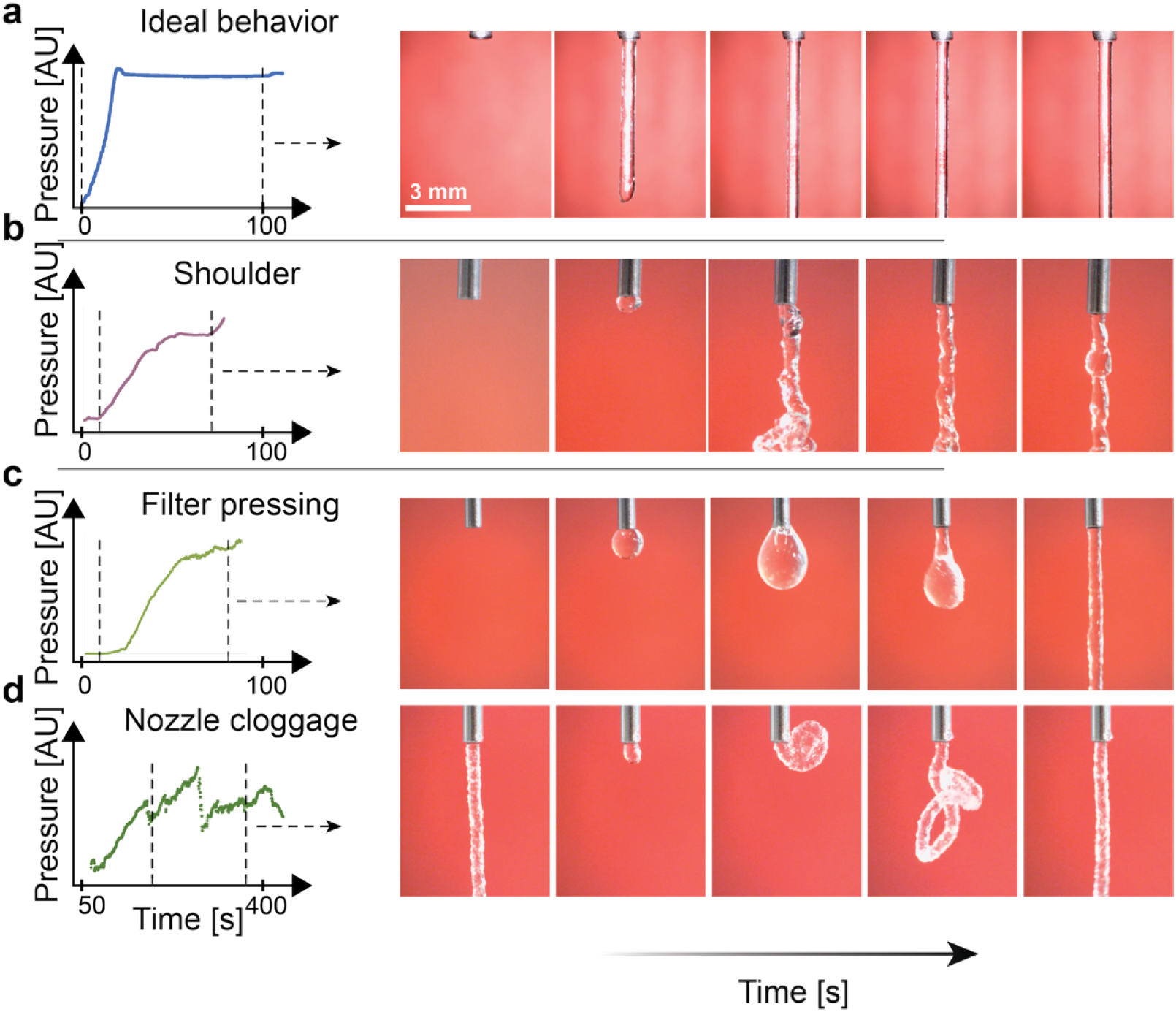
Extrusion phenomena captured using X-TRUDE across three hydrogel systems. (a-d) Pressure curves paired with corresponding microscopic images for each formulation. (a) Pluronic-25 exhibits stable, homogeneous extrusion with minimal temporal variation. (b) mGelMA-2.5 displays a pronounced shoulder feature and fragmented extrudate. gGelMA reveals (c) a filter-pressing regime, characterized by an initial low-pressure aqueous phase followed by a rapid pressure increase, as well as (d) intermittent nozzle clogging, reflected by rapid pressure spikes and transient pressure drops. All experiments were carried out at room temperature (21 °C). AU indicates arbitrary units.

X-TRUDE-based pressure patterns revealed two characteristic phases in the extrusion process across all formulations. The first phase, termed the “onset phase,” represents the initial transient period characterized by a rapid pressure rise as the material begins to fill the nozzle. The second phase, the “process phase,” corresponds to the steady extrusion regime, during which pressure values exhibit time-dependent fluctuations and intermittent instabilities reflecting evolving hydrogel-geometry interactions. The complete set of pressure-derived extrusion features extracted from the pressure traces, along with their identification criteria, is summarized in **Table S1**.

To illustrate how these phases manifest across three representative classes of hydrogels, we first examined Pluronic-25 as a model system known to exhibit stable, homogeneous extrusion behavior.^[57,58]^ For Pluronic-25, the pressure increased sharply (with mean slope of 1.14 Pa s⁻¹) as the hydrogel progressively filled the empty nozzle during the onset phase, ultimately stabilizing at a steady value (799.4 ± 6.2 Pa) once Pluronic-25 exited the capillary tip (Figure 4a). After reaching this equilibrium point, the pressure remained nearly constant throughout the entire duration of the extrusion process (t = 104 min), confirming that Pluronic undergoes a highly uniform and temporally stable process phase.

In contrast to Pluronic’s steady and uniform extrusion profile, mGelMA-2.5 exhibited a pronounced “shoulder effect,” characterized by an initial pressure rise followed by a transient plateau (Figure 4b). During the first 60 s, the pressure rose from 0 to ∼1500 Pa, plateauing at t = 60 s for roughly 20 s. This transient plateau corresponds to material exit from the nozzle tip as confirmed using microscopic observation. Notably, mGelMA-2.5 resulted in a clear delineation between initial and later stages of extrusion compared to Pluronic. Subsequent pressure increase beyond the shoulder effect (starting at t ∼ 80s) found to reflect the transition of extrusion process to a stage where pressure built-up balances resistance to flow. Microscopic observations revealed that mGelMA-2.5 exhibited notable non-uniform extrusion, reflected by a fragmented extrudate structure (Figure 4b) with irregular edge morphology, consistent with reports from glass-capillary extrusion.^[58]^

To broaden the scope of our extrusion analysis beyond homogeneous and monolithic hydrogels, we next investigated gGelMA as a jammed granular formulation that represents an emerging hydrogel class.^[59]^ In contrast to mGelMA, gGelMA exhibited additional transient flow phenomena. Most notably, gGelMA displayed filter-pressing phenomenon,^[54]^ which is a pressure-induced phase-separation process involving preferential expulsion of interstitial liquid phase from a material prior to extrusion of the solid-like (granular) phase. X-TRUDE results demonstrated that filter-pressing phenomenon divided the extrusion process into two distinct phases: initially, the aqueous phase was the primary component to be expelled, reflected by near-zero (2.3 ± 1.2 mbar) pressure readings (Figure 4c). After ∼30 s, a rapid increase (slope = 14.2 mbar s^-1^) in pressure was observed, corresponding to the combined-phase extrusion following the partial depletion of aqueous phase. To further investigate the pressure patterns resulting from this phenomenon, we examined pressure data from a long (38.1 mm) and a short (∼1 mm) cylindrical nozzle simultaneously. During the initial stage of filter pressing, the syringe equipped with the longer nozzle exhibited near-zero pressure, indicating minimal hydrodynamic resistance, whereas the shorter capillary registered higher pressure. This counterintuitive result suggests that the expelled fluid was of low viscosity—primarily aqueous phase—rather than the granular hydrogel phase. Under viscoelastic material extrusion conditions, pressure drop is expected to directly scale with capillary length due to cumulative viscous losses.^[18]^ Therefore, the reversal of this expected pressure-length relationship provides strong evidence of phase separation during the initial stages of extrusion. Once the hydrogel-rich phase began to extrude, the pressure measured using the longer capillary increased and surpassed that of the shorter one, consistent with theoretical predictions for single-phase, viscoelastic hydrogel flow through cylindrical conduits.^[10,18]^ It should be noted that the filter-pressing inherently alters the local composition of the extruded material, leading to spatiotemporal gradients in rheological properties that deviate from those of the originally prepared formulation (**Figure S19**).

Beyond phase separation, structurally heterogeneous materials are more prone to exhibiting additional failure modes during extrusion, most notably nozzle clogging. X-TRUDE measurements using gGelMA revealed a characteristic pressure signature for clogging events, consisting of rapid pressure spikes followed by abrupt drops and subsequent recovery (Figure 4d). These transient signatures reflect transient obstruction at the nozzle entrance or within the converging region, where local accumulation of aggregates or densely packed particulates temporarily impedes flow.

Although these experiments focused on gGelMA, the underlying clogging behavior is consistent with established frameworks of confined soft-matter flow, where local jamming or particle accumulation can arise when discrete microgels or particulates migrate, compact, or orient unfavorably within constricted geometries.^[60]^ This mechanistic linkage is particularly relevant for emerging classes of hydrogels that incorporate microgels, particles, or other modular building blocks.^[59,61]^ By resolving clogging signatures with high temporal sensitivity, X-TRUDE can enable early identification of flow obstruction and offers a route to gain mechanistic insight and rationally optimize formulations and nozzle architecture to ensure consistent extrusion outcomes.

### 2.3. High-fidelity hydrogel extrusion requires spatiotemporal thermal control

While the mechanical shearing condition of a hydrogel is a primary determinant of extrusion performance, temperature acts as an equally critical yet often uncontrolled variable.^[16,62,63]^ This insight is particularly relevant since many commonly used hydrogel systems, such as gelatin and poly(N-isopropylacrylamide),^[64,65]^ exhibit thermosensitive behavior. Accordingly, even small temperature variations might significantly alter their physico-mechanical properties, thereby profoundly affecting extrusion behavior. Nevertheless, conventional rheological methods are typically conducted under uniform temperature conditions,^[54]^ and thus fail to capture these coupled thermal-flow dynamics. To address this gap, we leveraged the X-TRUDE system to quantify how controlled thermal conditions—representative of realistic extrusion environments (e.g., during 3D bioprinting)—govern the pressure evolution and filament formation of a representative thermosensitive hydrogel (i.e., mGelMA-2.5). To this end, we examined three thermal conditions that reflect typical extrusion application scenarios,^[3,41,54,66]^ linking fundamental gelation transitions with realistic extrusion conditions: (i) isothermal baseline condition (21 °C), (ii) non-isothermal condition (30 °C syringe / 21 °C chamber), and (iii) elevated isothermal condition (30 °C).

Under the isothermal baseline condition, both the material and the chamber were equilibrated to room temperature (21 °C) prior to extrusion. Under these conditions, mGelMA-2.5 exhibited non-uniform filament formation, as evidenced by local fluctuations in the pressure data. This non-uniformity was further corroborated by microscopic imaging, showing inconsistencies in filament morphology throughout the extrusion process (**Figure 5a**).

**Figure 5.**
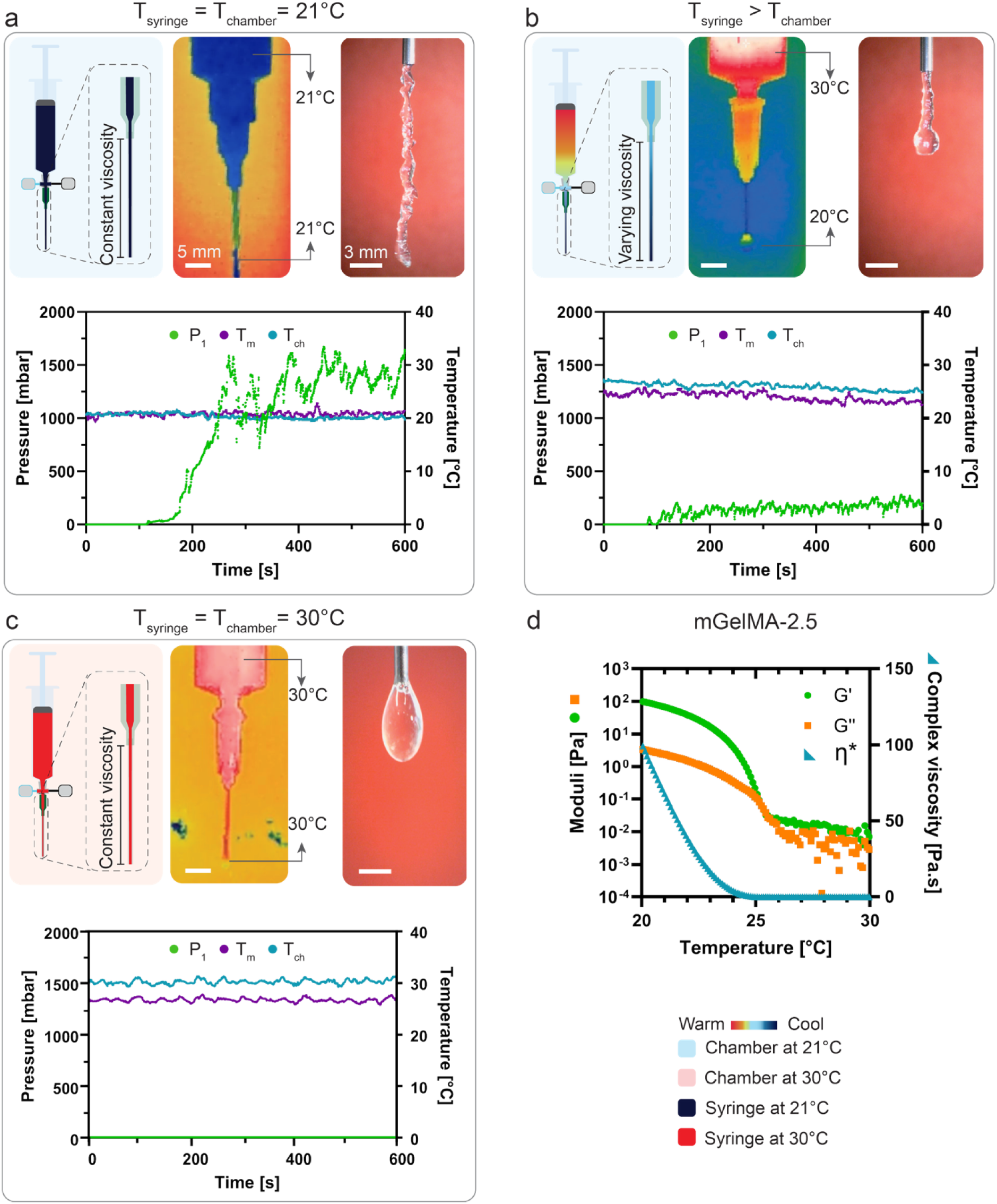
Effects of variation in local and environmental temperature on hydrogel extrusion behavior measured using the X-TRUDE system. (a-c) Schematic illustrations, infrared (IR) thermal images, microscope images of the extrudate, and corresponding material temperature (T_m_), chamber temperature (T_ch_), and pressure (P_1_) curves under three different temperature conditions: (a) isothermal baseline condition at 21 °C for both syringe and chamber, (b) non-isothermal condition with syringe at 30 °C and chamber at 21 °C and (c) elevated isothermal condition with both syringe and chamber equilibrated at 30 °C. (d) Complex viscosity and viscoelastic response of mGelMA-2.5 obtained during a continuous temperature sweep from 30 °C to 20 °C.

Next, we evaluated the non-isothermal condition by preconditioning the material to 30°C within the syringe while maintaining the chamber at room temperature (21 °C). This configuration generated a temperature difference of ∼9 °C between the material and printing chamber, resulting in a transient extrusion behavior. This setup emulates realistic extrusion scenarios frequently encountered in 3D (bio)printing,^[67,68]^ where pre-warmed GelMA encounters a cooler nozzle and/or chamber environment, inducing dynamic sol-gel transitions that frequently lead to extrusion irregularities.

Under this condition, the material initially extruded with minimal resistance, as evidenced by low pressure readings (7.8 ± 0.8 mbar). However, as extrusion progressed, the pressure steadily increased (reaching 166.8 ± 3 mbar at 10 min), indicating time-dependent stiffening of the hydrogel (Figure 5b). Infrared (IR) thermal imaging revealed that as the material left the syringe and contacted the cooler capillary walls (initially ∼25 °C), it rapidly cooled from 30 °C to ∼26 °C, while simultaneously transferring heat to the sensor adapter (i.e., custom-made adapter for sensor integration into the material extrusion path) and the capillary. Over time, the adapter gradually warmed up, while the mGelMA in the syringe lost heat to the surrounding environment through the syringe wall (∼28 °C to ∼26 °C in 2 min, **Figure S20**). This process established two superimposed thermal gradients: (i) a radial gradient between the syringe and the environment, and (ii) a vertical gradient along the extrusion path from the syringe reservoir to the capillary tip (**Figure S21-S23**).

These temperature gradients influence extrusion through two coupled mechanisms. (i) Thermo-rheological effect: mGelMA’s local viscoelasticity varies with temperature, generating a spatially heterogeneous flow. As evidenced by rheological temperature sweeps, temperature change from 30 °C to 25 °C results in a ∼17 and ∼26-fold increase in G’ and G’’, respectively, (Figure 5d). As a result temperature gradients along the flow path, the material exhibits progressively increasing viscoelastic resistance during extrusion, resulting in pressure build-up and increasingly irregular filament morphology. (ii) Thermal history effect: mGelMA exhibits a time-dependent viscoelastic response when it is exposed to a temperature change, particularly near its sol–gel transition temperature.^[3,69]^ The characteristic timescale of this network response is on the order of several seconds,^[69,70]^ which can overlap with the material’s residence time inside the nozzle depending on the applied flow rate. Under these conditions, the hydrogel experiences spatiotemporal evolution of its viscoelastic properties during flow as the material traverses the nozzle. As a result, extrusion behavior is governed not only by the local temperature and applied extrusion force, but also by the thermal history that controls the degree of network formation at each point along the flow path. This complex interplay between gelation kinetics, thermal history, and flow rate produces a non-equilibrium extrusion response that cannot be captured by conventional rheological measurements conducted under uniform temperature conditions.

Finally, we subjected the system to the elevated isothermal condition under which both the material and the chamber were maintained at 30 °C for one hour prior to extrusion to ensure thermal equilibrium. Extrusion was caried out at the same temperature. Under this condition, mGelMA-2.5 resided above its sol-gel transition window, and therefore behaved as a low-viscosity, weakly elastic solution. During extrusion, the material cannot sustain any filament structure and instead breaks into droplets immediately upon exiting the nozzle (Figure 5c). Correspondingly, the extrusion pressure remained nearly constant at 200 ± 15 mbar, which is consistent with flow response governed almost entirely by viscous dissipation, rather than any appreciable elastic or yield-like contributions. This regime provides an important upper bound in the thermal landscape of GelMA extrusion. When the material is fully in the sol state, elastic network formation is negligible, and the hydrogel cannot support its own weight once deposited. As a result, filament integrity is lost and capillary breakup becomes the dominant instability mechanism. These observations highlight that extrusion uniformity is compromised not only by excessive cooling, as in the non-isothermal case, but also by excessive heating that suppresses gelation entirely. The elevated isothermal condition therefore complements the isothermal baseline and non-isothermal regimes by illustrating the full thermal span across which mGelMA transitions from fragmented flow, to transient stiffening, to complete loss of filament stability.

Overall, our findings underscore that the extrusion of thermosensitive hydrogels is governed by a coupled temperature-time-flow landscape rather than by material viscosity alone. Spatial temperature gradients create immediate variations in viscoelasticity, while thermal history determines how rapidly a network rearranges as the material’s temperature changes inside the nozzle and/or post extrusion. When these processes evolve on the same timescale as the residence time in the nozzle, the resulting flow is fundamentally out of equilibrium, leading to more complex pressure evolution and filament morphologies that conventional isothermal rheology cannot predict. By recapitulating spatiotemporal temperature variations of extrusion processes, X-TRUDE directly quantifies these coupled mechanisms and identifies the thermal operating window in which stable filaments can be produced.

### 2.4. In situ pressure profiles correlate to filament morphology during extrusion

Qualitative assessment of the pressure profiles indicated that formulations exhibiting large, irregular fluctuations (e.g., mGelMA-2.5-21°C) produced highly unstable, buckled filaments, whereas formulations with smooth pressure evolution (e.g., Pluronic-25) yielded straight, uniform filaments (Figure 6a). This qualitative correspondence motivated a systematic, quantitative analysis of whether the characteristics of these fluctuations can predict filament morphology across compositions. To this end, we analyzed mGelMA across a range of polymer concentrations (i.e., 2.5, 5, and 10 wt%) and isothermal extrusion temperatures (i.e., 21 °C and 28 °C) using the X-TRUDE system.

**Figure 6.**
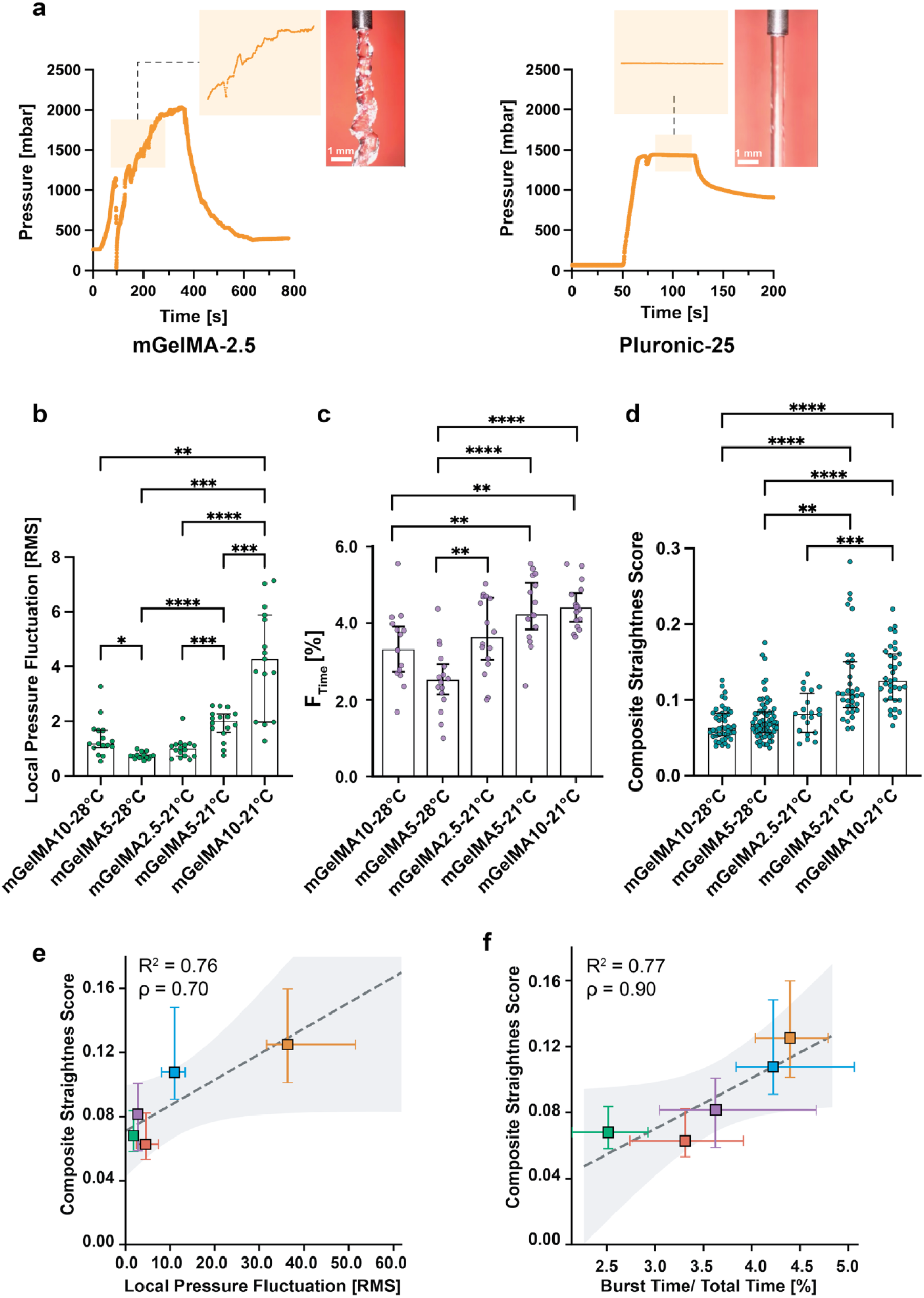
Analysis of filament morphology using pressure and visual data. a) Representative pressure curves, corresponding local pressure fluctuations, and microscopic images of extruded filaments for mGelMA-2.5 and Pluronic-25 formulations. (b-d) Effects of polymer concentration (2.5, 5, and 10 wt%) and extrusion temperature (21 °C and 28 °C) on filament morphology for mGelMA-2.5, mGelMA-5, and mGelMA-10 quantified by (b) local pressure fluctuation (LPF), (c) fraction of time spent in burst states (F_time_), and (d) overall filament uniformity expressed as a composite straightness score. (e-f) Correlation between LPF amplitude and composite straightness score (e), and F_time_ and composite straightness score (f) quantified using Spearman method. (b-d) *, p < 0.05; **, p < 0.01; ***, p < 0.001; ****, p < 0.0001; statistical significance for (b–d) was determined by Kruskal-Wallis H-tests followed by Mann-Whitney U tests with Bonferroni adjustment for multiple comparisons. Data in panels (b-f) are presented as median and interquartile range (IQR).

Closer examination of the raw pressure profiles revealed two distinct temporal components: (i) a low-frequency global trend, reflecting gradual syringe pressurization and flow stabilization within the syringe-nozzle assembly, and (ii) a superimposed relatively high-frequency local component, originating from rapid, local pressure perturbations associated with microscale flow instabilities in the material. To isolate these high-frequency local fluctuations, we detrended each pressure profile for its low frequency behavior, enabling quantification of rapid fluctuations independent of low-frequency drift. Subsequently, we first studied the magnitude of the detrended fluctuations using a local pressure fluctuation (LPF) metric, defined as the absolute deviation of the detrended pressure from its baseline (zero) value. LPF increased markedly with mGelMA concentration and decreasing temperature (Figure 6b). While amplitude-based metrics capture the magnitude of pressure fluctuations, they do not describe how long the flow remains in an unstable regime. Accordingly, we computed the fraction of the total experimental time during which bursts event occurred (F_time_). This analysis revealed that stiffer or colder mGelMA formulations exhibited not only larger fluctuation amplitudes but also significantly longer periods of sustained instability (Figure 6c), whereas softer formulations displayed only brief, transient events.

To compare these pressure-derived trends with extrusion outcomes, we measured filament straightness from optical images using a composite morphological score. The morphology measurements followed the same ordering across compositions as the LPF values (Figure 6d), indicating conditions with larger pressure fluctuations consistently yielded fewer straight filaments.

Samples exhibiting low LPF values, such as mGelMA-2.5-21°C and mGelMA-10-28°C, generated straight and uniform filaments. These low-amplitude, short-timescale fluctuations indicate that the material experiences only minor variations in local flow resistance as it traverses the nozzle. In practical terms, this behavior reflects a stable balance between viscous dissipation and elastic deformation, enabling continuous and uniform filament formation. In contrast, conditions associated with large-amplitude or unstable pressure fluctuations, such as mGelMA-10-21°C, consistently produced irregular, fragmented, or tortuous filaments (Figure 6d). High LPF values signify rapid and repeated perturbations in local stress, which are commonly associated with transient jamming–unjamming events, localized yielding, or abrupt elastic recoil within the hydrogel network.^[58]^ These microscopic instabilities interrupt the continuity of flow at the nozzle exit, resulting in visible morphological defects. The strong correspondence between LPF amplitude and filament irregularity further supports the mechanistic link between short-timescale viscoelastic dynamics and macroscopic extrusion quality.

Finally, we assessed whether pressure-derived features could directly predict filament morphology. Despite the small number of conditions tested, both LPF and F_time_ showed monotonic correlations with the composite straightness score (Figure 6e-f), with LPF and F_time_ are associated with 76% and 77% of the variance in this limited dataset, respectively. Nonetheless, the slightly stronger correlation between F_time_ and filament morphology (ρ = 0.90, compared with ρ = 0.70 for LPF) indicates that the temporal persistence of instability may be more influential for determining filament morphology than fluctuation magnitude alone.

The use of pressure fluctuations as a proxy for flow instability is supported by polymer-extrusion studies, where characteristic pressure oscillations have been linked to the onset of jerky stop-and-go flow (i.e., stick–slip), the development of a rough, ribbed surface on the extrudate (i.e., sharkskin), and severe distortion or breakup of the extrudate (i.e., melt fracture). Signal processing of pressure traces has therefore been proposed for on-line quality monitoring of extrusion processes.^[5,71–73]^ While we have demonstrated here for mGelMA and Pluronic F-127, this framework is in principle applicable to other viscoelastic compositions.

These findings establish that filament morphology can be inferred directly from pressure data alone, without the need for image-based evaluation. This insight represents a major step toward real-time, closed-loop process monitoring in extrusion-based processes. By enabling non-invasive prediction of extrusion performance, this method provides a mechanistic framework for optimizing material formulation and extrusion conditions, ultimately enhancing reproducibility and control in biofabrication workflows.

## 3. Conclusions

Reliable hydrogel extrusion is governed by the coupled interplay of intrinsic materials properties and the extrinsic processing conditions. Conventional RR, despite serving as the field’s primary characterization method, probes only idealized shear fields and uniform isothermal environments. Accordingly, available methodologies fail to capture the heterogeneous stress states, thermal gradients, and temporal flow instabilities that govern extrusion performance in processes such as extrusion-based 3D (bio)printing.

The X-TRUDE system presented in this work closes this gap through enabling a process-informed framework. By integrating in situ pressure sensing, spatiotemporal thermal control, and controlled geometric constraints, X-TRUDE captures the combined shear and extensional stress contribution, thermal gradients, and temporal flow transitions that dictate extrusion performance. This capability extends beyond the viscosity measurements provided by conventional capillary rheology systems, enabling the detection of extrusion-specific features such as onset effects, process effects, filter-pressing, and clogging.

Our comparative analyses across different material formulations revealed that shear-thinning behavior alone does not predict extrudability. Instead, extrusion outcomes correlate with the combined effects of extensional resistance, temperature-dependent viscoelasticity, flow-path geometry, and formulation heterogeneities. In many extrusion systems, hydrogels are preconditioned at temperatures different from that of the nozzle or surrounding environment,^[3,41,54,66]^ yet this mismatch is rarely monitored or controlled. X-TRUDE testing demonstrated that such non-isothermal conditions induce spatial heterogeneity in viscoelasticity, and that the resulting thermal history significantly modifies flow behavior over the extrusion timescale. Across material classes, formulations express distinct extrusion failure modes. For instance, granular hydrogels are more prone to phase-separation phenomena such as filter-pressing. Analysis of pressure fluctuations, particularly through local pressure fluctuation (LPF), mechanistically links these transient instabilities to the resulting filament morphology, providing a generalizable sensory metric for predicting extrusion performance across diverse material systems.

Positioning X-TRUDE within the broader context of extrusion technologies requires distinguishing extrudability from printability. Extrudability refers to stable and continuous material delivery through a nozzle, whereas printability additionally encompasses post-extrusion behaviors such as filament shape retention, substrate interaction, structural relaxation, and self-healing.^[41,46,51,74]^ By resolving material behavior under realistic thermal and geometric constraints, X-TRUDE establishes the mechanistic basis for predictive extrudability assessment, which forms the first and essential component of overall printability. Nonetheless, the implications of X-TRUDE extend beyond 3D (bio)printing. Processes involving the extrusion of viscoelastic soft materials, including minimally invasive therapeutics, cosmetic fillers, soft-matter manufacturing, and food processing face similar challenges of transient flow regimes, thermal gradients, and geometry-dependent stresses. By enabling early identification of extrusion failure modes and revealing the mechanistic origins of flow heterogeneity, X-TRUDE has the potential to shorten development cycles, reduce material waste, and guide rational formulation design across a broad range of technologies.

Overall, this work establishes X-TRUDE as a mechanistic and predictive framework for evaluating hydrogel extrusion under conditions that match real processing environments. By bridging the longstanding gap between conventional rheological characterization and actual extrusion performance, X-TRUDE advances the field towards more reproducible, reliable, and process-informed engineering of hydrogel-based systems. Accordingly, the integration of X-TRUDE framework into material development pipelines has the potential to significantly enhance the fidelity, robustness, and translational reliability of soft-matter extrusion technologies in biofabrication and beyond.

## 4. Experimental

*A brief overview of key experimental procedures is provided below. Comprehensive protocols, system specifications, and additional methodological details are available in the Supporting Information*.

### Design and assembly of X-TRUDE system

Briefly, the system consists of a temperature-controlled extrusion chamber fabricated from laser-cut poly(methyl methacrylate) (PMMA; Plexiglas) panels and an extrusion module constructed from aluminum struts, machined profiles, and 3D printed components produced using a Formlabs 3B+ printer (Formlabs, USA). Real-time pressure measurements were acquired using a high-sensitivity inline pressure sensor (ELVH series, Amphenol All Sensors, USA) positioned directly upstream of the nozzle, and in situ temperature was monitored through a thermistor (GA10K3MRBD1, Polyamide NTC Thermistor, TE Connectivity, USA) positioned upstream of the nozzle. All hardware components were operated through an Arduino Uno WiFi microcontroller (Arduino, Italy) that communicated with a custom Python interface for synchronized control of extrusion parameters, environmental temperature, and continuous acquisition of sensor data.

### Preparation of hydrogel formulations

Pluronic F-127 (Pluronic-25): Pluronic F-127 powder (Sigma-Aldrich) was dissolved in 1× phosphate-buffered saline (PBS, Sigma-Aldrich) to a final concentration of 25 wt%. The solution was stirred at 600 rpm for 4 h in an ice–water bath until complete dissolution. Solutions were then transferred into 10 mL syringes and stored at 4 °C until use.

Monolithic GelMA (mGelMA): mGelMA solutions (2.5, 5, and 10 wt%) were prepared by dissolving X-Pure GelMA 160P60 (160 kDa, 60% degree of methacrylation, Rousselot BV, Netherlands) in PBS at 70 °C for 30 min under magnetic stirring (600 rpm). After dissolution, solutions were transferred into 10 mL syringes and physically crosslinked at room temperature (∼21 °C) for 30 min, then stored at 4 °C until further use. The second GelMA batch with identical specifications (mGelMA-B2) was prepared using the same procedure and used immediately after physical crosslinking at room temperature.

Granular GelMA (gGelMA): Extrusion-fragmented GelMA (granular hydrogel) was prepared in multiple steps. First, 5 wt% GelMA powder (first batch GelMA) was dissolved in PBS at 70 °C for 30 min and mixed with a separately prepared 12.5 wt% Lithium phenyl-2,4,6-trimethylbenzoylphosphinate (LAP, Sigma-Aldrich) stock solution at 40 °C to achieve a final LAP concentration of 0.25 wt%. The mixture was then transferred to 20 mL syringes and physically crosslinked at room temperature (∼21 °C) for 30 minutes and then at 4 °C for 10 min, and subsequently photocrosslinked under UV light (365 nm, 15 mW cm⁻², 120 s, flipped midway). The crosslinked hydrogel was then extruded sequentially through decreasing needle sizes (18G to 30G), washed with PBS, and centrifuged (10,000 RCF) to obtain a jammed microgel suspension. The suspension was then transferred to 10 mL syringes and used immediately afterwards.

### Rotational Rheology

Oscillatory and steady shear measurements were carried out to characterize rheological behavior of the materials behavior using rotational rheometry (RR) and to compare these results with those obtained from X-TRUDE. Measurements were conducted on a AR2000 Advanced Rheometer (TA Instruments, USA) equipped with a 40 mm parallel-plate geometry. The frequency and shear-rate ranges were matched between RR and X-TRUDE experiments to enable direct comparison of the resulting data.

### X-TRUDE Analysis of Hydrogel Formulations

After mounting hydrogel-loaded syringes onto the syringe holders of X-TRUDE system, the plunger was positioned in contact with syringe pistons. Extrusion tests were conducted with an applied linear extrusion rate in descending order: 15, 10, 5, and 1 mm s^-1^, corresponding to flow rates of 498.70, 332.50, 166.25, 33.20 µL min^-1^, respectively, using a 12 mm long (18G) nozzle. Each extrusion step was maintained at these constant shear rates until a steady-state pressure plateau was reached. Real-time pressure values were automatically recorded by a custom Python code for further analysis. To accurately mimic the initial phase of common extrusion-based applications, both the sensor adapter and capillaries were kept empty prior to start of extrusion. This experimental approach ensured that recorded pressure values began at zero mbar.

### Microscopic video acquisition

Filament morphology was monitored using a Dino-Lite digital microscope (AM7915MZT, Dino-Lite, Taiwan) with red background contrast for better visualization. The microscope was positioned at an approximate distance of 25 cm from the capillary and kept at this fixed distance for all the acquisitions. Video recording began 10 s before extrusion initiation for temporal synchronization with pressure data.

### Filament Morphology Analysis

Filament morphology was quantitatively assessed using custom Python workflow employing AI-based segmentation with pre-trained deep learning models. Although an intensity-based multi-method consensus edge detection approach (gradient, adaptive thresholding, zero-crossing, and K-means clustering) was initially implemented, it did not reliably detect filament boundaries across all imaging conditions and was therefore not used for final analysis. Raw filament images were converted to grayscale and subjected to Gaussian blur processing using a 5×5 kernel to reduce noise. Canny edge detection was applied with optimized low and high thresholds (6 and 32, respectively) to identify filament boundaries. Binary masks were generated using a scanline fill algorithm that connected the first and last detected edge pixels in each row, followed by vertical gap filling (maximum gap: 3 pixels) to ensure continuous filament boundaries.

AI-based segmentation outputs were used to extract filament boundaries, from which six morphological parameters were computed: edge root mean square error (RMSE), centerline straightness RMSE, width coefficient of variation (CV), width standard deviation, normalized maximum edge deviation, and a composite straightness score (mean of edge RMSE, centerline RMSE, and width CV). All deviation-based metrics were normalized by mean filament width to enable comparison across different filament diameters.

### Filament local fluctuation quantification

Pressure fluctuation analysis was performed on real-time extrusion data sampled at ∼19 Hz for all mGelMA formulations. Local pressure fluctuations were quantified by first removing slow baseline drift using a discrete wavelet–based adaptive detrending approach (db4, level-8 approximation), which isolates rapid, transient pressure events while preserving their temporal structure. Bursts were defined as high-amplitude events exceeding twice the standard deviation of the detrended signal (|P_d_(t)| > 2σ). For each burst, duration and peak amplitude were extracted, and two quantitative metrics were calculated: (i) the fraction of total extrusion time spent in the burst state (F_time_), and (ii) the root-mean-square (RMS) magnitude of the detrended pressure signal, representing the overall fluctuation intensity. Detailed algorithms and parameter choices are provided in the Supporting Information.

### Infrared thermal data acquisition

An infrared camera (HIKMICRO Mini2Plus; HIKMICROTECH, China) was employed to capture thermal images and videos of entire syringe-capillary assembly surface at 25 Hz. The camera was positioned at a fixed distance of 20 cm from the surface of the syringe and was focused on the syringe-capillary assembly surface. The emissivity index (ɛ) was set to 0.96, an established value for the polymeric material of the syringe assembly. This setting ensured accurate conversion of emitted thermal radiation into the true surface temperature of the assembly. Due to the opacity of the plastic in the long-wave infrared spectrum, the thermal images captured the surface temperature profile of the syringe, which serves as an indicator of the internal temperature of the hydrogel and its thermal transfer to the environment. The camera’s contrast mode was adjusted to “rainbow” mode to enhance the syringe visibility against the background for later analysis.

### Statistical analysis

Statistical analyses were performed in Python (SciPy, statsmodels, pandas). Data were assessed for normality using the Shapiro–Wilk test and for homogeneity of variance using Levene’s test. Due to significant departures from normality and unequal variances, non-parametric methods were employed for primary inferential analyses. Differences across GelMA composition–temperature groups were evaluated using the Kruskal-Wallis H-test. Following a significant omnibus result, post-hoc pairwise comparisons were performed using Mann–Whitney U tests with Bonferroni-adjusted significance levels to control the family-wise Type I error.

The relationship between extrusion pressure fluctuations and filament straightness was assessed using Spearman rank correlation (ρ) as the primary inferential statistic, supplemented by Pearson correlation for comparison. Linear regression with 95% confidence intervals was used for visualization. Data are presented as median and interquartile range (IQR). All tests were two-tailed with significance set at α=0.05 unless otherwise noted.

## Acknowledgements

F.S. and M.D. would like to thank Dr. Felix Hol for providing access to the laser cutter and Dr. Felix Evers for his assistance in laser cutting process. This work was funded by Hypatia Grant (BoneChipPredict) from Radboud University Medical Center and VICI grant from The Netherlands Organization for Scientific Research (NWO, grant no. 17835).

## Conflict of Interest

The authors would like to disclose a submitted patent application related to the X-TRUDE system reported in this manuscript. International Patent Application filed with the European Patent Office (EPO) on 20 November 2025 by Radboud University Medical Center (Application No. EP25217512.0; Priority date: 20 November 2025). Inventors: F.S, P.B, M.D.

## Data Availability Statement

((include as appropriate, including link to repository))

Received: ((will be filled in by the editorial staff)) Revised: ((will be filled in by the editorial staff)) Published online: ((will be filled in by the editorial staff))

## Supporting Information

Supporting Information is available from the Wiley Online Library or from the author.

## Notes

### Competing Interest Statement

The authors disclose a submitted patent application related to the X-TRUDE system described in this manuscript. International Patent Application filed with the European Patent Office (EPO) on 20 November 2025 by Radboud University Medical Center (Application No. EP25217512.0; Priority date: 20 November 2025). Inventors: F.S., P.B., and M.D.

