## Supporting Information for "X-TRUDE: A Process-Informed Framework for High-Fidelity Analysis of Hydrogel Extrusion"

### 1. X-TRUDE System Architecture

#### 1.1. Mechanical and Structural Components

The X-TRUDE system consists of an extrusion unit that integrates a mechanically driven syringe-based extruder, interchangeable syringes and capillaries, all incorporated within a thermally regulated chamber. All CAD models were developed using Autodesk Inventor Professional 2023 (Autodesk, USA).

##### *1.1.1. Mechanical Design of the Extrusion Unit*

The extrusion unit (**Figure S1**) is composed of an aluminum frame that provides mechanical rigidity. Two aluminum strut profiles (Bosch Rexroth, Germany), each measuring 6×3 cm, were assembled to create four slots accommodating an aluminum syringe mount and fixtures for the linear motion and actuating components.

The syringe holder was fabricated from an aluminum T-profile, while the stepper motor was mounted on an aluminum L-profile (MetaalShopper, Netherlands). To mount these profiles on the backbone struts, connection holes were machined using a vertical drill, and threads were tapped using a handheld threading tool (**Figure S2**).

A stepper motor (2.7 N m, 103H7 Series, Sanyo Denki, Japan) drove a lead screw (TR8×2, 8 mm diameter) through a shaft coupling to convert rotational motion to linear plunger displacement.

##### *1.1.2. 3D-Printed Components*

Key components were 3D printed using Black Material v4 resin and a Form 3B+ printer (Formlabs, USA) to ensure high precision in guiding the syringe plunger and ensuring accurate alignment of components. A custom 3D-printed holder with integrated V-slots was incorporated into the aluminum T-profile to secure syringes and prevent undesired movement during operation. This component included a threaded nut that, when fastened with a syringe clamp, firmly held syringes in position. The syringe plunger was guided by two linear bearings (LBBR 14, Ewellix, Sweden) mounted on parallel guide rods (14 mm diameter, Ewellix), ensuring smooth and precise motion. The guide rods were secured by 3D-printed top holder at top and laser-cut mounting points on the chamber floor (**Figure S3**).

*1.1.3. Chamber and Enclosure Fabrication*

The chamber dimensions were specified as 58 × 40 × 40 cm to accommodate syringes up to 20 mL capacity along with the longest capillaries while providing adequate space for convenient handling during operation. The chamber was constructed from 5 mm poly(methyl methacrylate) (PMMA; Plexiglas) sheets, which were cut according to design files generated by an open-source online software (https://boxes.boringplace.org/). A "simple box" template served as the foundation with dimensions of 40 × 40 × 58 cm (width × depth × height). The bottom edge utilized a "Stackable" format to promote efficient airflow to electronics. Design parameters included 5 mm sheet thickness and 0.14 mm burn correction, with default settings retained for remaining options. The finalized design was exported as a Scalable Vector Graphics (SVG) file and imported into Adobe AutoCAD 2023 (Adobe, California, USA) for modifications including openings for screw-nut assembly and openings for integrated component (i.e., heater, fan, rails) (**Figure S4**). Modified files were used to cut the PMMA sheets to their final form suing Q400 $CO_2$ laser cutter (Trotec, Austria). Detailed manufacturing drawings are available from the authors upon reasonable request.

#### 1.2. Sensor Integration

*1.2.1. Sensor Adapter*

An adapter was 3D printed with Clear Material V4 resin (Formlabs, USA) to enable sensor integration into the material extrusion path. The adapter featured four openings (**Figure S5**). Syringe and capillary connections established by Luer-lock connector system combining threaded and friction connections. Pressure and temperature sensor integration employed reversible sealing mechanisms. The pressure sensor connection was sealed using a silicon O-ring (RS PRO Silicon O-ring, RS Components, UK). The temperature sensor was secured to a threaded cap and sealed to the adapter using a silicon washer. The temperature sensor opening served a dual function, facilitating removal of air bubbles from the pressure sensor compartment. These openings were aligned to enable bubble removal, which was critical as trapped air introduces undesirable system compressibility, causing delays between material deformation and pressure sensing. To minimize pressure losses, the adapter's internal diameter was matched to those of the syringe tip and capillary (**Figure S5**).

##### *1.2.2. Pressure Sensor Assembly*

A pressure sensor assembly was developed to ensure stable and reliable electronic connections between the pressure sensors and the Arduino board, as well as to facilitate facile pressure sensor replacement. A printed circuit board (PCB) was designed to securely hold the pressure sensors while allowing connection via a USB-C port. The design of PCBs was outsourced to a third-party manufacturer (PCBWay, China). Additionally, a 3D printed enclosure was developed to protect the pressure sensor assembly from accidental damage (**Figure S6**).

##### *1.1.1. Temperature Sensor Assembly*

A high-accuracy thermistor (GA10K3MRBD1, Polyamide NTC Thermistor, TE Connectivity, USA) was employed to continuously monitor the temperature of the material flowing through the sensor adapter, ensuring accurate evaluation of the effects of local temperature variation on material properties. The thermistors were connected in series with 10 kΩ resistors to form a voltage-divider circuit **(Figure S7a)**. In this configuration, temperature-dependent changes in the thermistor's resistance produce corresponding changes in the output voltage. This voltage signal is recorded by the control electronics and converted to temperature using the thermistor's calibration curve.

The thermistors were attached to the adapter using a non-reversible adhesive applied to a thread-fitted cap (Figure S7b) ensuring stable placement. Additionally, to measure the chamber temperature, a thermocouple was positioned vertically inside the chamber within ~10 cm distance from the syringes and connected directly to the PID temperature controller. This thermocouple provided the feedback signal used by the PID controller to regulate the chamber temperature based on this physical point of reference.

#### 1.3. Electronics and Control Systems

##### *1.3.1. Power Distribution and Electrical Circuit*

The system operated from a 220 V, 50 Hz AC mains source supplied through an F-type plug. Power distribution was routed through a fused switch to all components (**Figure S8**). All AC wiring was performed using 4 mm² insulated cables. A 48 V, 7.32 A switching power supply (S8FS-C35048, Omron, Japan) provided power for the stepper motor and ensured sufficient current under maximum load. A dedicated 48 V, 2.2 A stepper motor driver (STR-2M, Applied Motion Systems, USA) with microstepping capability received control signals from the Arduino Uno Wi-Fi Rev3 board (Arduino, Italy). A secondary 12 V regulated power supply (TMP 15212C, TRACOPOWER, Switzerland) powered the Arduino board, a recirculation fan

(400 Series Axial Fan, ebmpapst, Germany), and a rapid cooling fan (8400 N Series Axial Fan, ebmpapst).

*1.3.2. Thermal Regulation System*

Temperature control was performed using a proportional–integral–derivative (PID) controller (E5CD-QX2A6M-002, Omron, Japan), connected to a Type K exposed-junction thermocouple (RS Pro) and a 400 W fan heater (Stego, USA). The heater power was selected based on heat-loss calculations for a sealed chamber to enable rapid heating and stable maintenance of 50 °C over prolonged operation. The PID controller regulated heater power via an analog-output-triggered relay. The thermocouple was positioned at the height of the lower syringe barrel to measure the temperature directly at the syringe location (Figure S8). A recirculation fan inside the chamber ensured uniform temperature distribution.

*1.3.3. Motor Driver and Motion Control Module*

Stepper motor actuation was achieved using the STR-2M motor driver powered at 48 V. The Arduino Uno Wi-Fi served as the motion controller, providing step and direction signals to the motor driver. Motor motion was controlled using the open-source AccelStepper library (https://www.airspayce.com/mikem/arduino/AccelStepper/), enabling smooth acceleration and deceleration during syringe extrusion. Maximum torque and microstepping configurations were programmed according to the manufacturer's specifications.

*1.3.4. Computer Interface and Data Acquisition*

The X-TRUDE system communicated with a host computer through a serial interface managed by the Arduino Uno Wi-Fi. The Arduino simultaneously controlled the stepper motor driver and acquired data from integrated pressure and temperature sensors. A custom Python application initiated the extrusion tests by sending motion commands to the Arduino. During operation, real-time sensor data were streamed back to Python, where they were logged and processed for analysis.

##### 1.4. System-Level Assembly of the X-TRUDE System

All structural components described in previous sections were assembled into the final X-TRUDE system. The chamber walls were interlocked using the designed press-fit joints, and internal mechanical components were mounted using the screw–nut fastening system. The extrusion rails and plunger assembly were aligned and secured to the back wall of the chamber to ensure smooth linear motion. Sensor modules, including the pressure and temperature ports, were positioned in their designated mounts, and the wiring for all sensors and actuators was routed through the chamber using cable guides (Natural Cable Tie Mount, Legrand) to prevent interference with moving parts. Once assembled, the chamber door was sealed to minimize heat loss, and the complete system was inspected for mechanical alignment and sensor functionality before calibration (**Figure S9**).

#### 2. System Calibration and Characterization

##### 2.1. Flow Rate Calibration

Flow rate calibration was performed at low-pressure and high-pressure environments.

###### *2.1.1. Low-pressure calibration*

Distilled water (density 1.0 g $mL^{-1}$) was used as the calibration fluid at room temperature (21 ± 1 °C). Target flow rates between 10 and 2000 µL $min^{-1}$ (**Table S2**) were imposed. For each flow rate, the effluent was collected in pre-weighed containers over a fixed time interval, and the mass was measured using a precision balance (Mettler Toledo; readability 0.001 g). The volumetric flow rate was calculated from the collected mass assuming water density $\rho = 1.0$ g $mL^{-1}$.

###### *2.1.2. High-pressure calibration*

The syringe outlet was connected to 10 mL plastic BD Plastipak™ syringes terminated with cylindrical stainless-steel dispensing tips (Nordson, USA) of different gauges (18G, 22G, and 25G; length 1 inch; internal diameters 0.84, 0.41, and 0.25 mm, respectively). A broad range of flow rates was applied (**Table S2**), and each flow rate was tested in triplicate for every needle gauge. The resulting data were used to calibrate the pressure sensor and validate the accuracy of flow rates under high-pressure.

To assess accuracy of flow rates both in short and long-time scales, experiments were carried out with two measurement regimes of "short" and "long" duration. To ensure a clearly measurable mass while limiting evaporation, target collection volumes of 10–200 µL (short) and 500–1000 µL (long) were selected. Each condition was measured in triplicate.

##### 2.2. Thermal Calibration

Thermal calibration of the thermocouples and thermistors was performed in a high-accuracy laboratory (Memmert). Each sensor was equilibrated for 15 min at each set temperature. Calibration proceeded in two phases:

###### *2.2.1. Air calibration*

Both the thermistors and the thermocouple were suspended in the oven air volume, and the recorded temperatures were compared to the oven set-point.

*2.2.2. Water calibration*

Both the thermistors and the thermocouple were immersed in a 50 mL beaker filled with tap water placed inside the oven. Sensor readings were compared to both the oven set-point and an additional reference mercury thermometer immersed in the same beaker.

Across the tested temperature range, thermocouples showed an accuracy within ± 0.5 °C, and thermistors within ± 0.1 °C, consistent with the manufacturers' tolerances.

**2.3. Viscosity Calibration**

The capillary-viscosity calibration of X-TRUDE was performed using S600 Viscosity Reference Standard (Paragon Scientific, UK), a Newtonian fluid with certified viscosity and negligible compressibility. Two 10 mL Fortuna Optima interchangeable glass syringes with Luer-lock fittings (Poulten & Graf GmbH, Germany) were filled with S600 and mounted in the chamber at 40 °C (reducing viscosity to ~0.4 Pa s to remain within the system pressure limits). Pressure drops were recorded across capillaries of different lengths and diameters using 4 bar pressure sensors (ELVH series, Amphenol All Sensors, USA) over the range of flow rates listed in **Table S3**. Apparent wall shear rate was calculated from pressure drop using Poiseuille's law:

$$\dot{\gamma} = \frac{4Q}{\pi R^3} \qquad \text{Equation 1}$$

where $Q$ is the volumetric flow rate, $\dot{\gamma}$ is the shear rate, and $R$ is the capillary radius.

Pressure losses due to flow convergence at the capillary entrance were corrected using Cogswell's method,[1] which uses pressure measurements from two capillaries with identical radii but different lengths (L/D ratio > 40). For Newtonian calibration fluids, only entrance effects were considered, assuming fully developed laminar flow, no slip at the wall, and atmospheric pressure at the capillary exit. The entrance pressure loss $\Delta P_{S1}$ was obtained as:

$$\Delta P_{S1} = (P_{1,S1} - P_{3,S1}) - (P_{1,S2} - P_{2,S2}) \qquad \text{Equation 2}$$

where $P_{1,S1}$ and $P_{3,S1}$ are pressures measured at points $P_1$ and $P_3$ of syringe 1(**Figure S10**) and the $P_{1,S2}$ and $P_{2,S2}$ are the pressures measured at points $P_1$ and $P_2$ of syringe 2. Since the pressure at tip of both capillaries was assumed to be atmospheric (i.e., 0) the equation 2 reduces to equation 3.

$$\Delta P_{S1} = (P_{1,S1}) - (P_{1,S2}) \qquad \text{Equation 3}$$

After correcting the recorded pressure values to account for entrance pressure losses, the true average wall shear stress in the capillary is determined using equation 4.

$$\tau = \frac{R\Delta P}{2L} \qquad \text{Equation 4}$$

where τ is the wall shear stress. *ΔP* is the pressure drop along the length of the capillary, *R* is the capillary inner radius, and *L* is the length of the capillary over which the pressure drop occurs. The viscosity of oil is then calculated using the equation 5.

$$\mu = \frac{\tau}{\dot{\gamma}} \qquad \text{Equation 5}$$

A calibration curve comparing $\mu$ values measured with X-TRUDE to the certified viscosity values of S600 Viscosity Reference Standard is shown in **Figure S11**.

**2.4. Syringe and System Compressibility Characterization**

System compressibility—a combined effect of syringe and material compressibility—introduced a lag in pressure response to step strain. Since materials could freely flow through the nozzle after step strain cessation, system compressibility manifested as material relaxation through pressure decay over time. To quantify this pressure decay, a modified version of a previously described method was used.[2]

To isolate system from material compressibility, the syringe piston was stopped at three positions after initiating extrusion from a fully loaded syringe, with distances measured relative to the syringe barrel beginning. By plotting material relaxation time versus stopping position, material relaxation, compressibility, and system compliance were determined. Relaxation time was defined as the time required for pressure to reach 95% of its lowest plateau value after flow cessation.

To isolate system compliance, experiments were conducted using a negligibly compressible material (S600 Viscosity Reference Standard) in three syringes: a 20 mL BD Plastipak™ 3-Piece syringe (Franklin Lakes, New Jersey, USA), a 10 mL Fortuna Optima Interchangeable glass syringe, and a custom 3D-printed (using Clear Resin V4, Formlabs) syringe. For low flowrates, stopping positions were closer to the capillary to minimize extrapolation, while for high flow rates, stopping positions were spaced further apart to allow adequate extrusion time for material response (**Figure S12a**).

Using a flow rate of 350 μL $min^{-1}$ through an 18G capillary, pressure values were monitored until equilibrium. At each stopping position, extrusion was abruptly halted and relaxation time determined. This process was repeated at three plunger positions for each syringe, with precise stopping positions determined using Equation 6.

$$L = \frac{V_i - (Q \times t)}{\pi \times R_i^2} \qquad \text{Equation 6}$$

where $V_i$ is the initial fluid volume inside the syringe, $Q$ is the volumetric flow rate, $t$ is the duration of applied flow, and $R_i$ is the internal radius of the syringe.

Plotting relaxation time at the three plunger positions provided values for both compliance and material relaxation. Extrapolating relaxation time to zero plunger position determined material relaxation. Since the selected materials exhibited negligible compressibility, observed compressibility resulted from syringe compliance (Figure S12b).

#### 3. Materials and Sample Preparation

##### 3.1. Hydrogel preparation

###### *3.1.1. Pluronic F-127*

Pluronic F-127 (Sigma-Aldrich) powder was dissolved in phosphate-buffered saline (PBS; Gibco) at a final concentration of 25 wt%. The powder was added gradually to cold PBS in an ice bath and gently stirred until complete dissolution (~4 h). The solution was transferred into 10 mL syringes and stored at 4 °C prior to use.

Pluronic F-127 exhibits a thermoresponsive sol–gel transition and forms a physically crosslinked hydrogel above ~20–25 °C due to micellization and micelle packing.[3] During all extrusion experiments, the material was handled at temperatures above the gelation temperature to ensure a stable hydrogel state within the syringe and capillary.

###### *3.1.2. Monolithic GelMA(mGelMA)*

mGelMA formulations with final concentrations of 2.5, 5, and 10 wt% were prepared following the CellInk protocol (CELLINK, Sweden). Briefly, GelMA powder was dissolved in PBS at 70 °C for 30 min in the dark (protection from ambient light with aluminum foil). The container was covered with Parafilm to minimize evaporation. No photoinitiator was added to these formulations. After dissolution, the warm solutions were transferred into 10 mL syringes and allowed to physically gel at room temperature (~21 °C). The syringes were then stored at 4 °C until extrusion.

###### *3.1.3. Granular GelMA (gGelMA)*

GelMA fragmentation took place using extrusion fragmentation method described previously.[4] Briefly, 5 wt% GelMA powder was dissolved in 80 (v/v)% of the final PBS volume and heated at 70 °C for 30 minutes. In parallel, a 12.5 wt% Lithium phenyl-2,4,6-trimethylbenzoylphosphinate (LAP) solution was prepared in the remaining 20 (v/v)% of the final PBS volume and heated for 30 minutes at 70 °C. The GelMA and LAP solutions were then combined at 40 °C to obtain the photocrosslinkable precursor.

The precursor solution was then transferred into 20 mL syringes and first physically gelled at 4 °C for 10 min, followed by chemical crosslinking by 365 nm UV illumination (15 mW $cm^{-2}$, 120 s), with the syringe flipped halfway through the exposure to ensure uniform crosslinking. The fully crosslinked bulk GelMA was then extruded sequentially through needles of decreasing size (18G to 30G) to generate a granular gel. The granules were washed twice with PBS by centrifugation at 8000 relative centrifugal force (RCF), followed by a final centrifugation at 10000 RCF to remove residual PBS. The granular suspension was finally transferred into 10 mL syringes and stored at 4 °C until use.

#### 4. X-TRUDE Experimental Protocols

##### 4.1. Extrusion Viscosity Measurement of Non-Newtonian Materials

The viscosity of non-Newtonian materials was determined using a procedure similar to that for Newtonian fluids, with additional post-processing to correct for their complex flow behavior. After loading two 10 mL Fortuna Optima Interchangeable glass syringes with the test material and mounting them onto the syringe holders, the X-TRUDE plunger was positioned to contact the pistons. Shear rates were applied stepwise from high to low, maintaining each step until a pressure plateau was reached. The pressure values were recorded for further analysis.

Using average pressure of last 30 data points (~1.5 second period), apparent viscosity was calculated using Poiseuille's law:

$$\mu = \frac{T_w}{\tau} = \frac{\pi \times R^4 \times \Delta P}{8 \times L \times Q} \qquad \text{Equation 7}$$

Where $T_w$ is wall shear stress, $\Delta p$ is the pressure difference between the two ends of capillary, $L$ is the length of capillary, $\mu$ is the dynamic viscosity, $Q$ is the volumetric flow rate, $R$ is the inner radius of capillary, and A is the cross-sectional area of capillary.

To accurately describe the flow behavior of shear-thinning hydrogels, two corrections were applied. The Weissenberg–Rabinowitsch (WR) correction[5] was employed to adjust the apparent shear rate, accounting for non-linear velocity profiles in the capillary:

$$\dot{\gamma}_{w} = \frac{3}{4} \times \dot{\gamma}_{ap} + \frac{1}{4} \times \tau_{W} \times \frac{d\dot{\gamma}_{ap}}{d\tau_{W}} \quad \text{Equation 8}$$

Where $\dot{\gamma}_w$ denotes true wall shear rate, $\dot{\gamma}_{ap}$ denotes apparent shear rate, and $\tau_W$ wall shear stress. This correction is particularly important at high shear rates and assumes a shear-thinning profile (**Figure S13**), which is a behavior commonly exhibited by hydrogels such as mGelMA and Pluronic.

Due to wall slip effects commonly seen in such hydrogels, the Mooney correction[6] was applied to estimate the sliding velocity at the capillary wall. This correction adjusts the apparent velocity profile derived under the no-slip boundary assumption:

$$y = \frac{v_g}{R} + f(\tau) \quad \text{Equation 9}$$

where $v_g$ is the sliding speed, $R$ is the capillary radius, and $\tau$ is the shear stress. For each shear stress level, a series of flow measurements was performed using capillaries with identical L/D ratios but different radii. Plotting shear rate against $R^{-1}$ yields the slip velocity from the slope.[6] The corrected viscosity values were then used to construct viscosity-shear rate curves for each material.

#### 4.2. Microscopy and Imaging

##### *4.2.1. Brightfield Imaging of Filament Morphology*

Filament morphology was captured using a Dino-Lite digital microscope (AM7915MZT, Dino-Lite, Taiwan) with red background contrast for enhanced visualization. Capillaries were emptied prior to each experiment to eliminate residual pressure effects. Recording began 10 seconds before extrusion initiation for temporal coordination with pressure data.

##### *4.2.2. Infrared Thermal Imaging*

An infrared camera (HIKMICRO Mini2Plus, HIKMICROTECH, China) was used to capture the temperature gradient across the material and syringe assembly during extrusion. The camera was positioned at a fixed distance of 20 cm from the surface of the syringe to ensure consistent measurement across all experiments. The emissivity index was set to 0.96, corresponding to the emissivity of water,[7] as hydrogels used in the experiments primarily consist of water. This setting allowed for accurate temperature measurement of the hydrogels. The camera's contrast mode was adjusted to “rainbow’’ to enhance the visibility of temperature variations within the material.

#### 5. Data Processing and Statistical Analysis

##### 5.1. Filament Morphology Quantification

Edge RMSE is root mean square error of edge deviations from fitted straight lines, calculated as:

$$RMSE_e = \frac{\sqrt{\frac{1}{n}\sum(d_{left,i}^2 - d_{right,i}^2)^2}}{w_{mean}} \quad \text{Equation 10}$$

where $d_{left,i}$ and $d_{right,i}$ are the deviations of left and right edges from their respective fitted lines at position $i$, and $w_{mean}$ is the mean filament width.

Centerline RMSE is deviation of the filament centerline from straightness:

$$RMSE_c = \frac{\sqrt{\frac{1}{n}\sum(c_i - \hat{c}_i)^2}}{w_{mean}} \quad \text{Equation 11}$$

where $c_i$ is the measured centerline position, $\hat{c}_i$ is the fitted straight-line value, and $n$ is the number of measurement points. Width Coefficient of Variation was defended as normalized width variability:

$$CV = \frac{\sigma_w}{w_{mean}} \quad \text{Equation 12}$$

where $\sigma_w$ is the standard deviation of filament width measurements.

The composite straightness score was calculated as the equally weighted average:

$$CSS = \frac{RMSE_e + RMSE_c + CV}{3} \quad \text{Equation 13}$$

All metrics were calculated using NumPy arrays for efficient computation. Linear regression (first-order polynomial fitting) was performed to establish reference straight lines for deviation calculations. Results were exported as CSV files for statistical analysis, with automated visualization overlays generated for quality control verification.

#### 5.2. Temperature Mapping from Infrared Recordings

To analyze temperature variations along the syringe, the recorded IR video was cropped to the size of syringe and then converted into a sequence of still frames using FFmpeg software (FFmpeg.org). Temperature mapping was performed by analyzing pixel values across multiple frames, corresponding to selected locations along the syringe.

A custom Python script was used to extract temperature data from these images. The first frame of the sequence was loaded as the reference image to specify four points along the syringe. As outlined in **Figure S21**, two of these points were selected along the capillary, one at the tip of capillary (P1) and the other at the base of capillary (P2) to ensure capturing temperature variance across the flow path. The other two points were selected within the syringe one closer to the outlet of syringe (P3) and the last one closer to the outer surface of the syringe but away from the syringe outlet (P4). These points were manually selected by the user through a graphical interface. To ensure consistent and comparable results across different experiments, the coordinates of these points were kept constants for all the IR images. The user was then prompted by the Python script to input the minimum and maximum temperatures corresponding to the color scale of the thermal images. These values were used to map pixel intensity (which lies between 0–255) to actual temperature values via linear interpolation. For each image in the sequence, the pixel intensity at the selected coordinates was extracted and converted into a temperature value. This procedure was repeated across all frames, producing a time series of temperatures for each selected point.

#### 5.3. Thermal Preconditioning and Analysis of Spatiotemporal Thermal Effects on mGelMA-2.5 Extrusion Behavior

For all the experimental groups ((i) isothermal baseline condition (21 °C), (ii) non-isothermal condition (30 °C syringe / 21 °C chamber), and (iii) elevated isothermal condition (30 °C)), mGelMA-2.5 was maintained at room temperature overnight before the experiment. For the elevated isothermal condition, after overnight preconditioning at room temperature, an additional two-hour conditioning at 30 °C was carried out in the X-TRUDE chamber prior to the initiation of extrusion tests. For the non-isothermal condition, mGelMA-2.5 was preconditioned at 30 °C for two hours and transported into the X-TRUDE chamber (set at 21 ± 1 °C) and start extrusion immediately. For each condition, extrusion tests were recorded using the IR camera as described in Section 4.2.2. For each recording, temperature time series were extracted at points P1–P4 along the syringe–nozzle–filament path (Figure S21).

For each point, the mean temperature over the steady-state extrusion period (last 60 s) and the temporal standard deviation were computed. In addition, the temperature difference between the syringe body (P4) and the nozzle/filament region (P1–P2) was used as a measure of axial thermal gradient. These metrics were compared across conditions to assess how preconditioning and chamber temperature affected (i) the uniformity of the temperature field and (ii) the resulting pressure profiles and filament morphology.

##### 5.4. Local Pressure Fluctuation (LPF) Analysis

###### *5.4.1. Adaptive Detrending Using Wavelet Transform*

To isolate local pressure fluctuations from slow baseline drift, wavelet-based adaptive detrending was applied using the discrete wavelet transform (DWT). As the pressure signals were non-stationary, this method allowed to preserve the temporal characteristics of pressure bursts while removing non-stationary baseline variations. Baseline detrending was performed using a discrete Daubechies 4 (db4) wavelet transform following previously established approaches for separating slow drift from localized dynamic events in non-stationary time-series data.[8] The detrended pressure signal $P_d(t)$ was obtained by subtracting the slowly varying baseline signal $B(t)$ from the raw signal $P(t)$:

$$P_d(t) = P(t) - B(t) \qquad \text{Equation 14}$$

where $B(t)$ was reconstructed from the level-8 approximation coefficients of a db4 wavelet transform.

###### *5.4.2. Burst Detection and Threshold Determination*

A burst defined as any pressure region during which:

$$|P_d(t)| > 2\sigma \qquad \text{Equation 15}$$

where σ is the standard deviation of $P_d(t)$. This $2\sigma$ threshold corresponds to selecting the top few percent (~2.5%) of fluctuations in a normal distribution. Binary burst state $B(t) = 1$ if $P_d(t) > 2\sigma$, and $B(t) = 0$ otherwise. Threshold-based burst detection has been extensively used across disciplines—including neuronal spike-train analysis, EEG dynamics, and transcriptional bursting—to identify rare, high-amplitude or temporally clustered events in complex biological signals.[9,10] Despite its broad use for characterizing neuronal activity and stochastic gene-expression kinetics, burst analysis has not been applied to quantify filament morphology in extrusion-based printing systems. Here we adapt the burst-detection principles to extrusion dynamics for quantitative identification of abrupt morphological events along extruded filaments. Each burst was characterized by duration, peak amplitude, and integrated energy.

*5.4.3. Fraction of Time in Burst State ($F_{time}$)*

To capture the temporal persistence of instability, the fraction of total time spent in burst states was computed:

$$F_{time} = \frac{\sum_i \Delta t_{burst,i}}{t_{total}} \qquad \text{Equation 16}$$

Where $\Delta t_{burst,i}$ is the duration of burst and $t_{total}$ is the total extrusion time.

*5.4.4. Root Mean Square Fluctuation Magnitude Calculation*

The root mean square (RMS) of the detrended pressure signal quantified the overall magnitude of pressure fluctuations:

$$RMS = \sqrt{\frac{1}{N}\sum P_d(t)^2} \qquad \text{Equation 17}$$

where N is the number of time points. For each experimental condition, median RMS and interquartile range (IQR) were calculated across all replicates. Median and IQR were used rather than mean and standard deviation due to the presence of occasional outlier measurements.

#### 5.5. Statistical Analysis

*5.5.1. Non-Parametric Statistics*

Given the non-normal distribution of the pressure fluctuation metrics and the moderate sample sizes, non-parametric statistical methods were employed for all primary inferential analyses. Data are presented as the Median, a robust measure of central tendency, accompanied by the IQR, defined as the 25$^{th}$ to 75$^{th}$ percentile range, which serves as a robust measure of data spread. Individual data points were plotted alongside these summary statistics to fully visualize the experimental distributions.

Differences across multiple experimental conditions (mGelMA concentration × temperature combinations) were assessed using the Kruskal-Wallis H-test, a non-parametric omnibus test. The Kruskal-Wallis H-statistic was calculated as:

$$H = \left(\frac{12}{N(N+1)}\right) \times \sum_i \left(\frac{R_i^2}{n_i}\right) - 3(N+1) \qquad \text{Equation 18}$$

where $N$ is the total sample size, $n_i$ is the sample size for condition $i$, and $R_i$ is the sum of ranks for condition $i$.

Following a statistically significant Kruskal-Wallis result, Mann-Whitney U tests were performed for non-parametric pairwise comparisons between conditions for post-hoc analysis. To maintain a family-wise alpha of 0.05, a Bonferroni correction was applied to all pairwise p-values.

*5.5.2. Correlation Analysis*

*Spearman Rank Correlation (ρ):* The Spearman Rank Correlation coefficient ($\rho$) was used as the primary statistic to assess monotonic association between pressure fluctuation metrics (RMS, $F_{time}$). This method is robust to non-normal distributions, reduced sensitivity to outliers, and requires no assumption of linearity in the functional relationship. The coefficient $\rho$ was calculated as:

$$\rho = 1 - \frac{(6 \times \sum d_i^2)}{(n(n^2 - 1)})$$ Equation 19

where $d_i$ is the difference between ranks for each paired observation. Spearman correlation was selected for its robustness to non-normal distributions and ability to detect monotonic relationships.

*Pearson Correlation (r):* The Pearson Correlation coefficient ($r$) was also calculated and reported for comparison, providing a parametric measure to quantify the strength of linear association. Correlations were computed using condition-level medians (N = 5 conditions: mGelMA-10-28°C, mGelMA-10-21°C, mGelMA-5-28°C, mGelMA-5-21°C, mGelMA-2.5-21°C) with two-tailed p-values reported at $\alpha = 0.05$.

The coefficient r was calculated as:

$$r = \frac{\sum[(x_i - \bar{x})(y_i - \bar{y})]}{\sqrt{\sum(x_i - \bar{x})^2 \times \sum(y_i - \bar{y})^2}}$$ Equation 20

The Coefficient of Determination ($R^2 = r^2$) was reported to represent the proportion of variance in morphology explained by the pressure fluctuation metric for the linear fit.

*5.5.3. Linear Regression and Confidence Intervals*

Linear regression models were fit for visualization using ordinary least squares:

$$\text{Morphology} = \beta_0 + \beta_1 \times \text{Pressure\_Metric} + \varepsilon$$ Equation 21

where $\beta_0$ is the intercept, $\beta_1$ is the slope, and $\varepsilon$ represents the residual error.

*5.5.4. Model Fitting and Confidence Interval*

The model was fit using Ordinary Least Squares (OLS) regression implemented via NumPy (polyfit function, degree=1). The 95% confidence interval for the regression line was calculated as:

$$CI = t_{(\alpha/2,n-2)} \times SE \times \sqrt{\frac{1}{n} + \frac{(x - \bar{x})^2}{\sum(x_i - \bar{x})^2}} \qquad \text{Equation 22}$$

where n is the number of data points, $t_{(\alpha/2,n-2)}$ is the critical t-value for 95% confidence with n-2 degrees of freedom, SE is the standard error of residuals, and $\bar{x}$ is the mean of the predictor variable. Given the limited number of conditions (N = 5), linear regression was primarily used for visualization and hypothesis generation. The small sample size limits the ability to conclusively distinguish between linear and non-linear functional forms. Consequently, the Spearman correlation, which makes no assumptions about the functional form, remains the primary inferential statistic.

##### 5.6. Calculation of average wall shear rates for 3D bioprinting applications

To determine average wall shear rates representative of bioprinting conditions, we drew on published extrusion parameters compiled in the public 3D bioprinting database curated by the Center for Engineering Complex Tissues (https://cect.umd.edu/database). Initially, all available bioprinting conditions (n = 531) employing cylindrical nozzles were identified. Conditions lacking clear specifications of nozzle diameter or printing speed (n = 60) were excluded from further analysis. For the remaining entries, the publications in which the internal nozzle diameters reported in Birmingham gauge units (n = 169), these diameter values were converted to metric units (i.e., mm) using the standard conversion tables (Nordson.com and Hamiltoncompany.com).

Maximum wall shear rate ($\dot{\gamma}_{wall}$) was determined based on the Hagen–Poiseuille flow model according to the following equations:

$$Q = VA \qquad \text{Equation 23}$$

$$\dot{\gamma}_{wall} = \frac{4Q}{60\pi r^3} \qquad \text{Equation 24}$$

where $Q$ is the volumetric flow rate, $V$ is the linear printing speed, A is the cross-sectional area of the nozzle, and r is the nozzle radius. Extrusion speed was assumed to be equal to the printing speed, a condition consistent with the majority of experimental setups in bioprinting field.[11] In this calculation, the assumption was that the extrusion material is a Newtonian fluid, making the calculated shear rate the highest possible rate during printing.

**6. Rotational Rheology**

Rotational rheology was carried out using an AR2000ex Advanced Rheometer (TA Instruments, USA). A 40 mm parallel plate steel geometry with a gap of 500 µm was used to conduct flow sweep (shear rate: 0.1 to 100 $s^{-1}$) and frequency sweep (1% strain amplitude, 0.1 to 100 rad $s^{-1}$ frequency) experiments, both at 21 °C. A 40 mm cone and plate steel geometry (2°) with a gap of 54 µm was also used for characterizing the shear-thinning behaviors of materials by flow sweep in forward (shear rate: 0.1 to 100 $s^{-1}$) and reverse (shear rate: 100 to 0.1 $s^{-1}$) direction at 21 °C. Finally, the 40 mm cone and plate steel geometry (2°) with a gap of 54 µm was used to conduct oscillatory temperature sweep (37 to 2 °C with 5 °C $min^{-1}$ cooling rate) and temperature ramp (37 to 2 °C with 60 seconds soak time) for characterizing the temperature responses of the material compositions.

**7. Supplementary Figures**

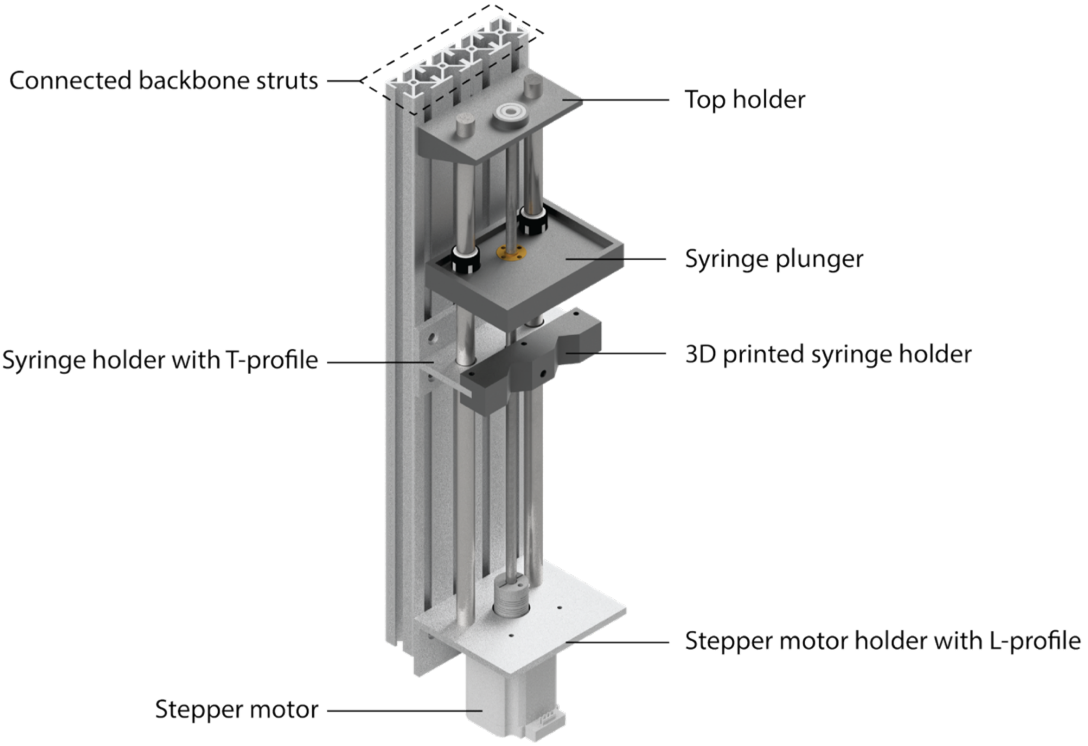

**Figure S1.** Perspective-view rendering of the extrusion unit assembly showing the aluminum backbone, machined, and 3D-printed structural components of the X-TRUDE system. The aluminum components include backbone struts, the stepper motor holder with L-profile, and the syringe holder base with the

T-profile. The 3D printed components include the syringe holder adapter mounted on the base T-profile, top mounting plate, and the syringe plunger.

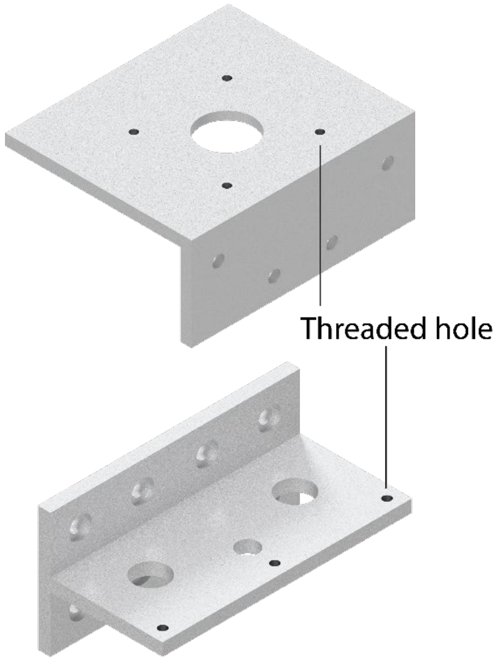

**Figure S2.** Machined aluminum components used in the extrusion unit. The upper bracket serves as the mounting interface for the stepper motor and includes threaded holes for secure attachment. The lower bracket serves as the mounting interface for syringes and contains through-holes and alignment features enabling connection to the alignment bars and structural rail system. All the holes were drilled using a manual vertical drill.

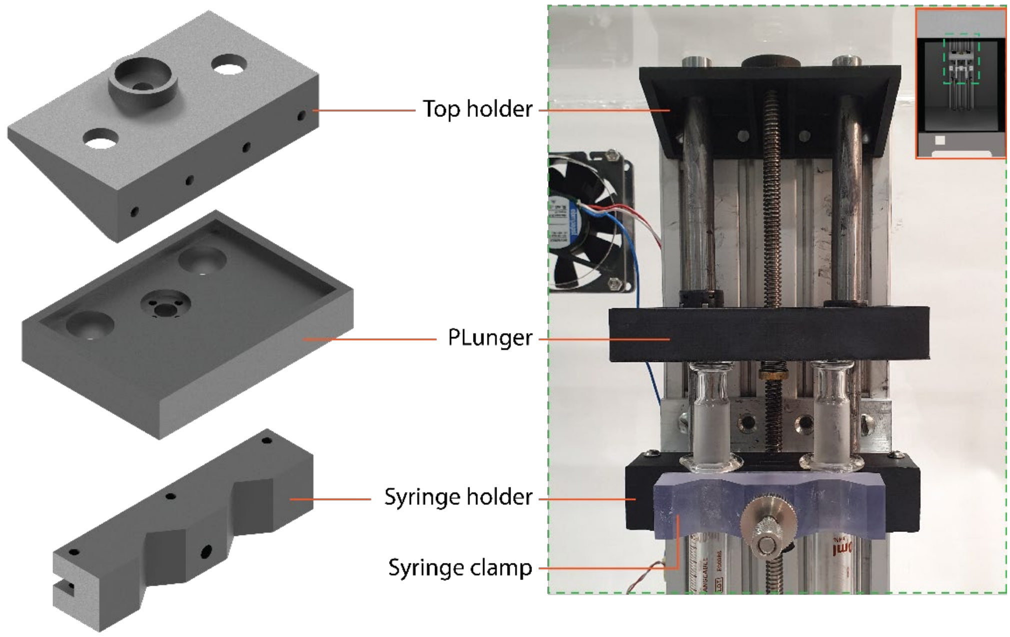

**Figure S3.** 3D-printed components of the extrusion assembly, including the top holder, plunger interface, syringe holder, and syringe clamp (left). These parts were assembled within the extrusion rail system, as shown in the photograph on the right. The highlighted region illustrates the installed 3D-printed components in their operational configuration.

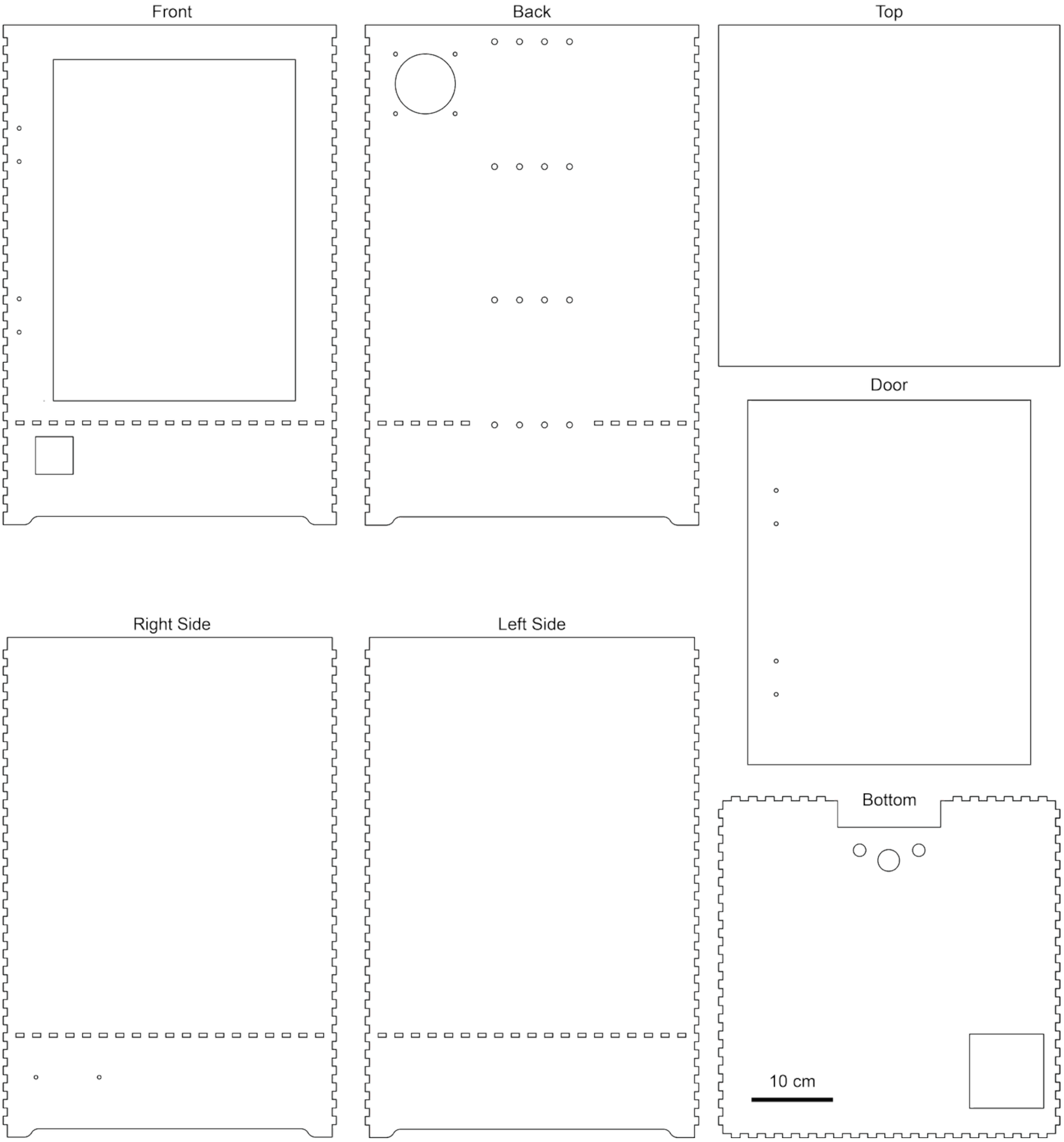

**Figure S4.** Schematic illustration of the individual chamber-wall components used for laser cutting from poly(methyl methacrylate) (PMMA; Plexiglas) panels. The layout includes the front, back, right side, left side, top, bottom, and door panels, each featuring predesigned openings for ventilation, cable routing, sensor ports, and mounting interfaces. Finger-joint edges are incorporated across all panels to enable precise alignment and tight assembly during construction.

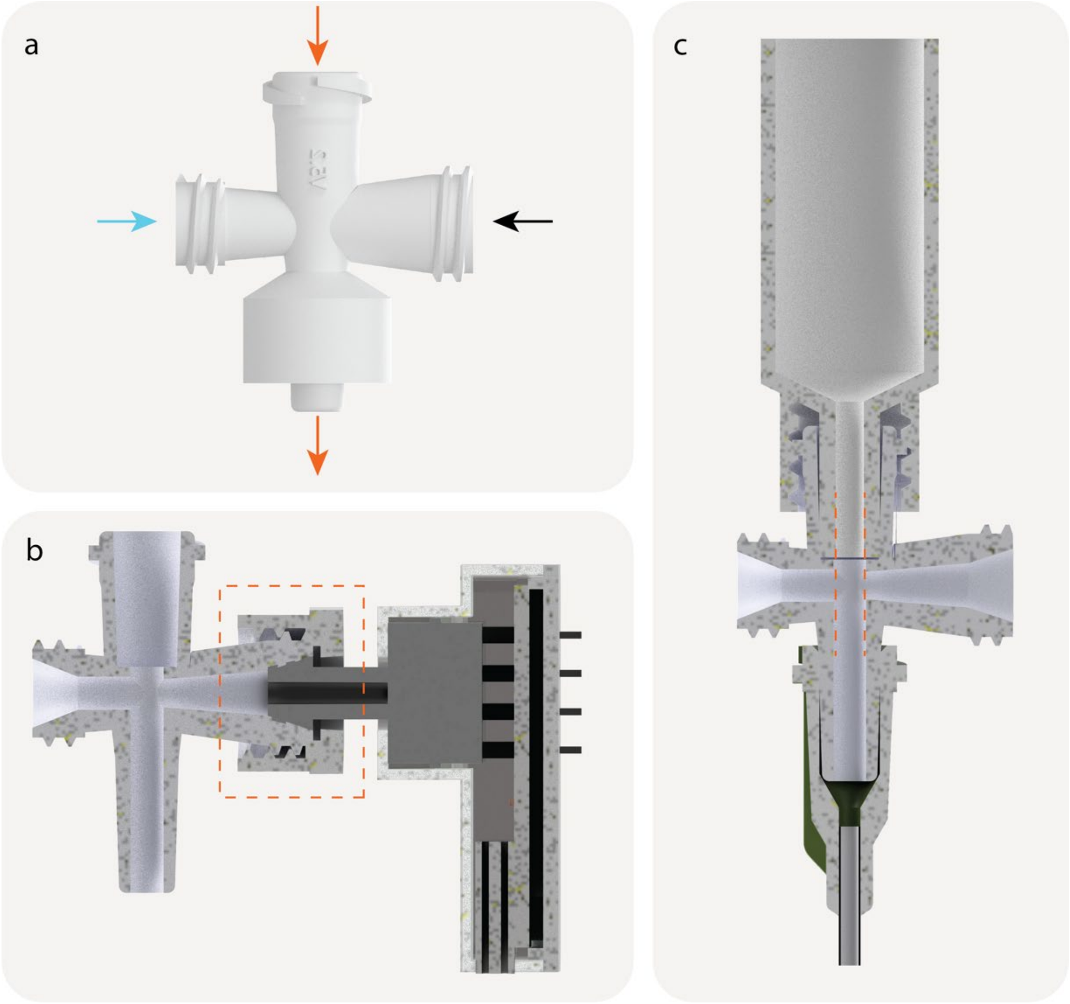

**Figure S5**. 3D rendering of the sensor adapter computer-aided design (CAD) and its assembly. a) The adapter with four connection points. Opening for syringe and nozzle connection and sensor integration are shown by orange and blue arrows, respectively. b) Cross-sectional view of the adapter and pressure sensor assembly. c) Cross-sectional view of syringe, the adapter, and nozzle assembly.

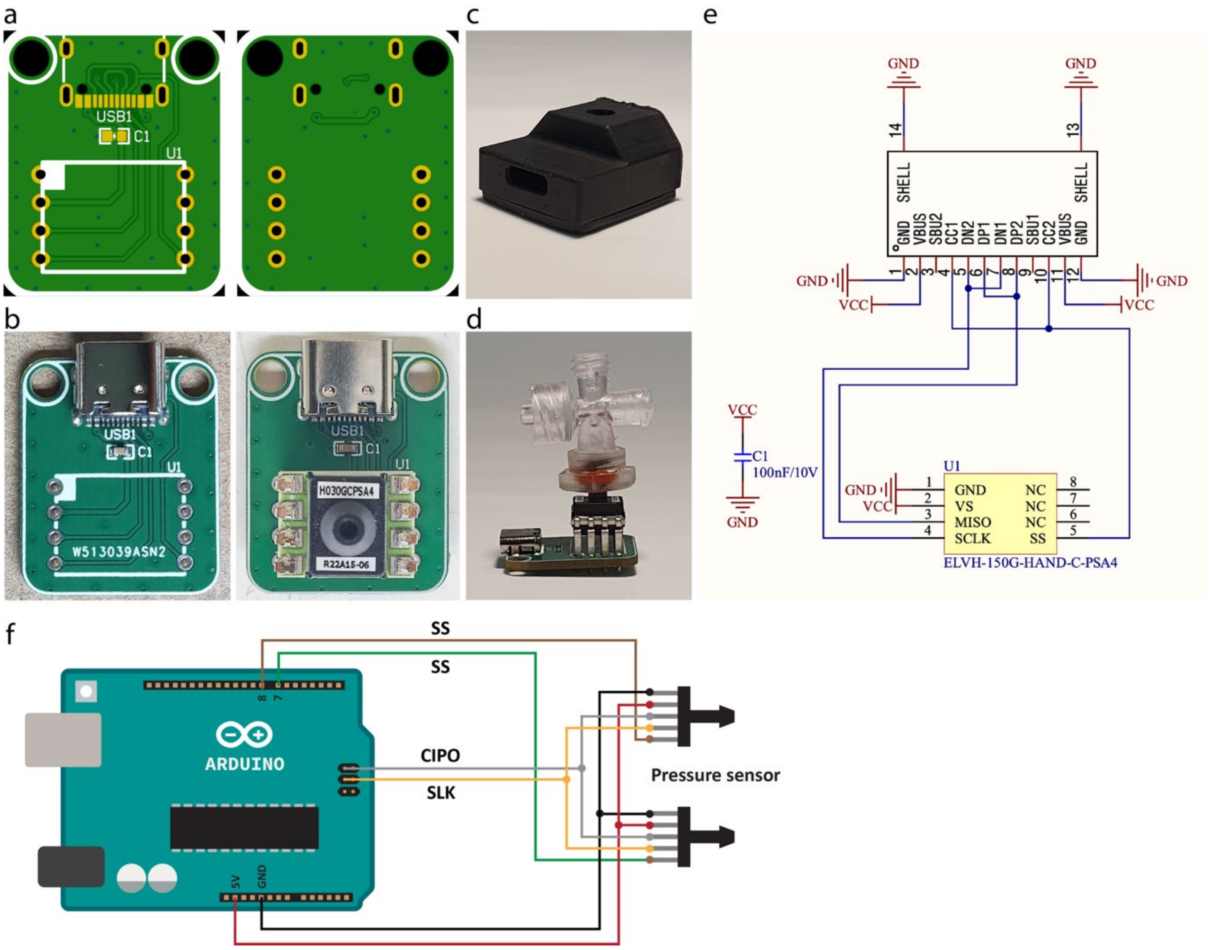

**Figure S6.** Pressure-sensor assembly and electrical connections. a) PCB layout of the custom-designed pressure-sensor board showing front and back layers. b) Photographs of the PCB before and after mounting the pressure-sensor chip. c) 3D-printed protective housing designed to enclose the assembled pressure-sensor module. d) Fully assembled pressure sensor attached to the custom sensor adapter. e) Schematic of the electrical connections between the USB-C interface and the pressure-sensor PCB. f) Wiring diagram illustrating the connection of two pressure sensors to the Arduino board using the Serial Peripheral Interface (SPI) protocol, with separate chip-select (SS) lines for each sensor, and shared CIPO and SCLK lines.

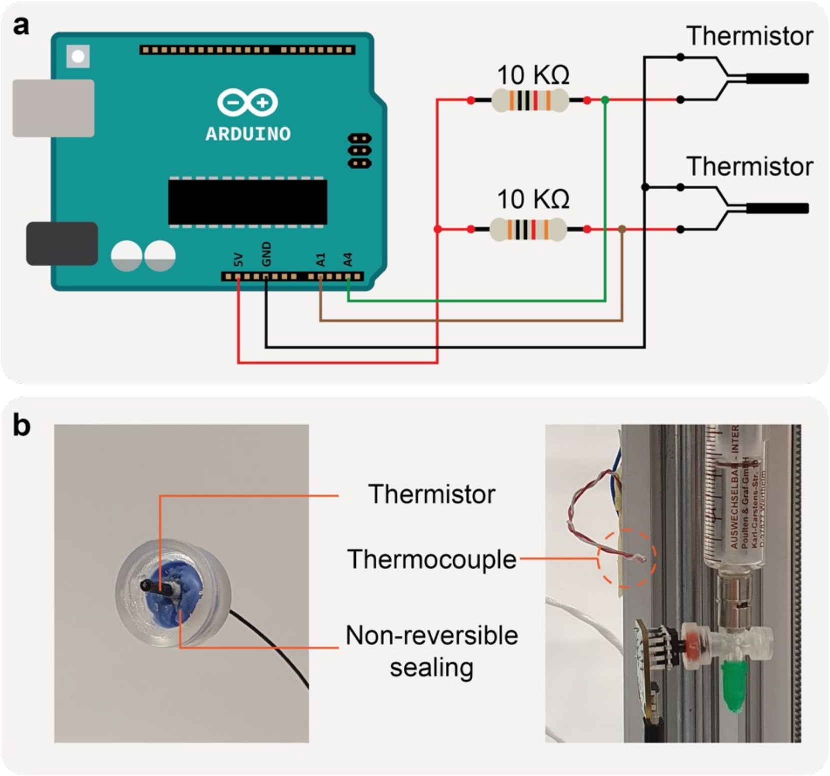

**Figure S7.** Electrical and mechanical integration of thermistors and the chamber thermocouple. a) Schematic of the thermistor circuits, each connected to the Arduino via a 10 kΩ voltage-divider configuration for analog temperature measurement. b) Photographs showing the mounted thermistors and the sealed attachment to the chamber cap using non-reversible sealing material, along with the placement of the thermocouple used for closed-loop chamber temperature monitoring.

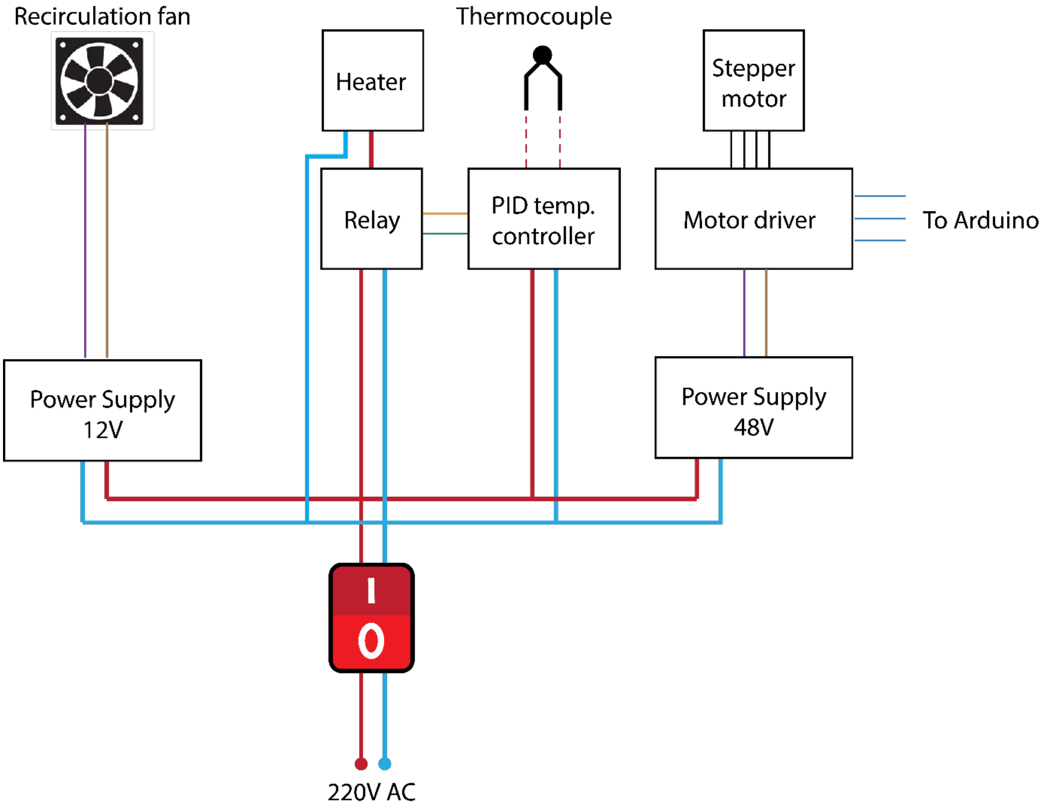

**Figure S8**. Schematic of the electrical power distribution for the X-TRUDE system. The 220 V AC mains supply feeds a master I/O switch, which distributes power to the 12 V and 48 V DC power supplies used for the recirculation fan and stepper-motor driver, respectively. The heater element is controlled through a relay driven by the PID temperature controller, which also receives input from the thermocouple. Low-voltage control signals, including the motor-driver inputs, are provided by the Arduino.

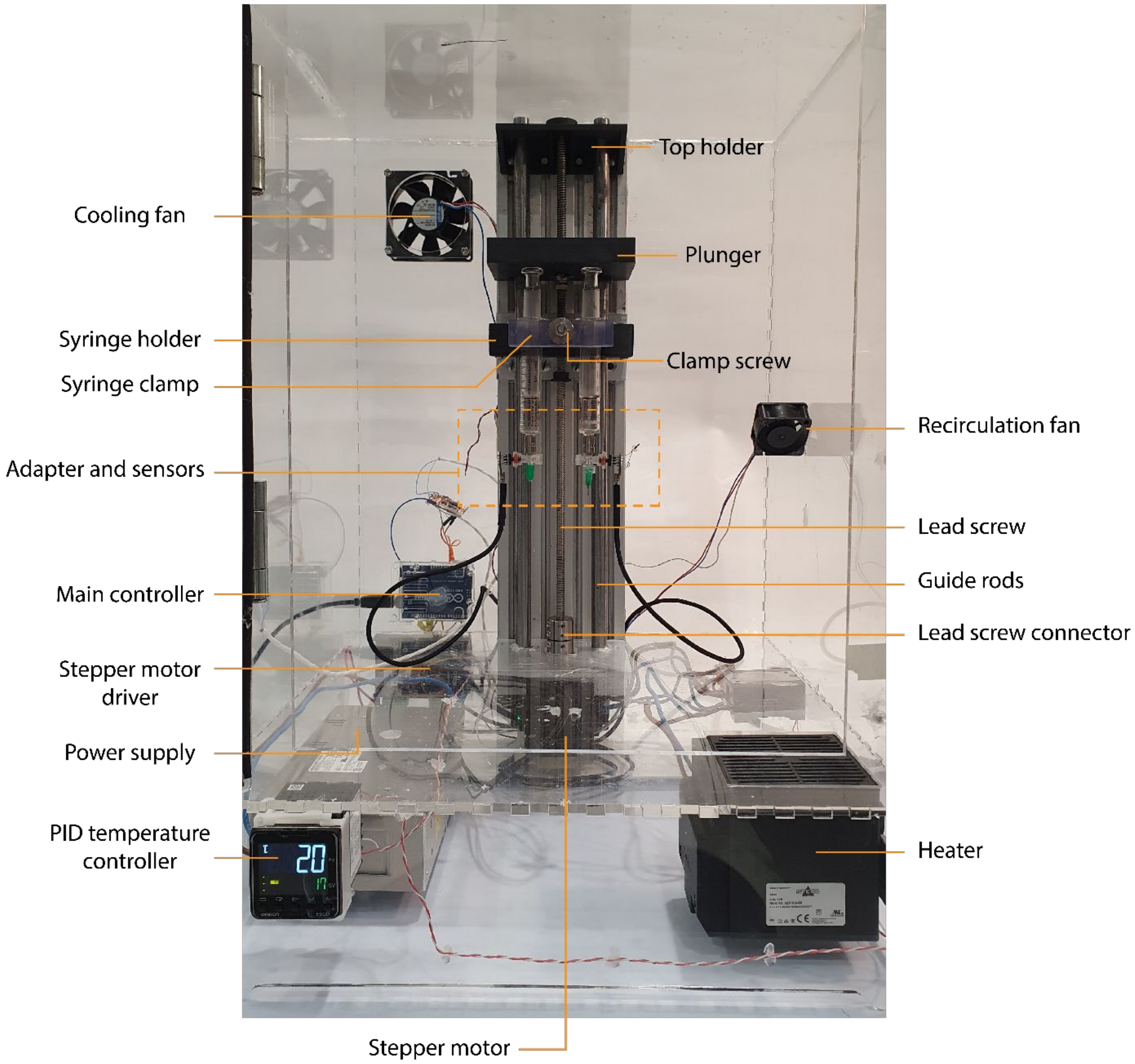

**Figure S9.** A digital photo of the fully assembled X-TRUDE system, showing the integration of the extrusion module (3D printed and machined aluminum components, stepper motor, and mechanical couplings), thermal-control components (heater, cooling and recirculation fans, PID controller), and the associated electronics (Arduino controller, motor driver, and power supplies).

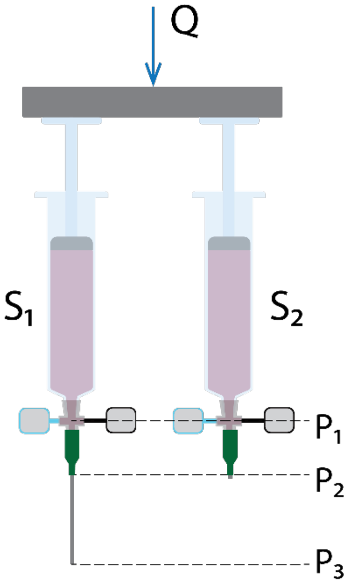

**Figure S10.** Illustration of the simultaneous extrusion of two syringes and the three different pressure acquisition points along the extrusion path. $S_1$ and $S_2$ refer to syringe 1 and syringe 2. $P_1$ is the pressure measured at the pressure-sensor location upstream of the capillary. $P_2$ is the pressure at the capillary inlet (equal to atmospheric pressure for syringe 2). $P_3$ is the pressure at the capillary tip, which is equal to atmospheric pressure. Q is the volumetric flow rate applied to both syringes assuming identical syringe dimensions.

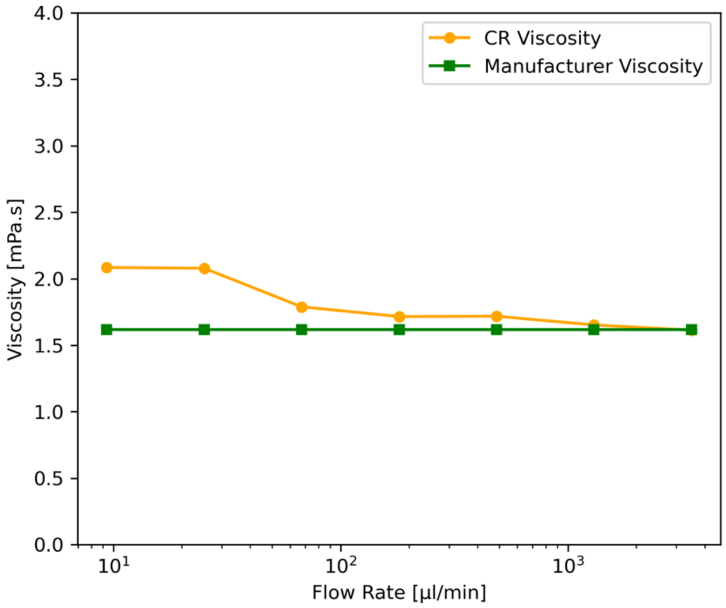

**Figure S11.** Calibration curves obtained using S600 Viscosity Reference Standard, comparing viscosities measured with the X-TRUDE to the manufacturer-provided viscosity values.

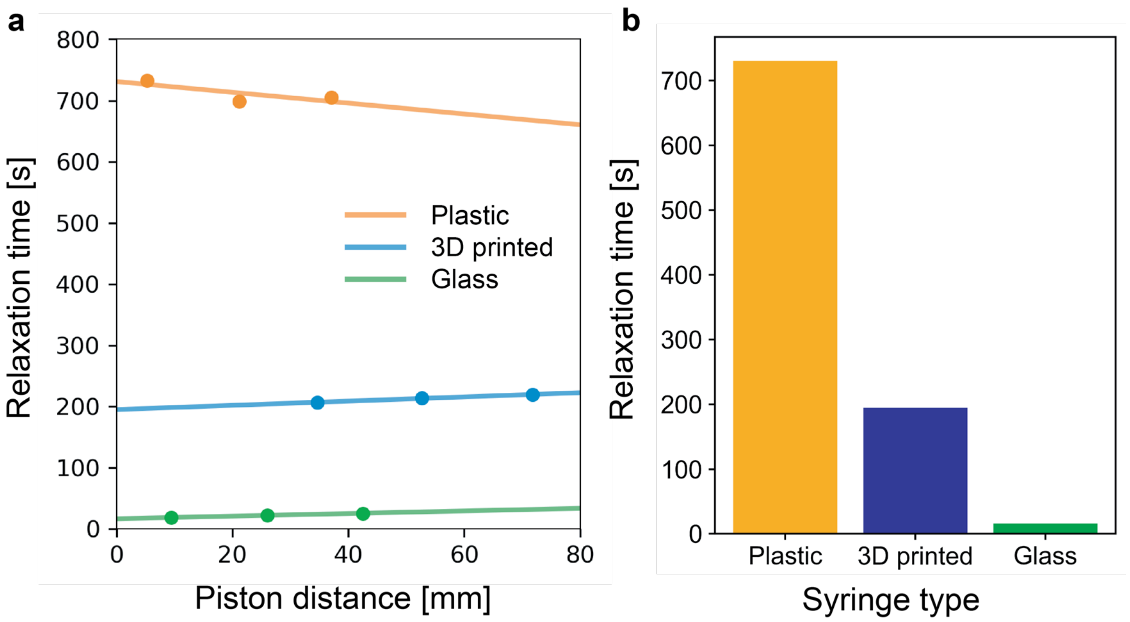

**Figure S12.** Effect of syringe material on syringe compressibility measured in the X-TRUDE system. a) Relaxation time as a function of piston tip stopping position measured from the syringe barrel beginning for a commercial plastic syringe, a 3D-printed syringe (Formlabs Clear V4 resin), and a commercial glass syringe, measured using S600 Viscosity Reference Standard at 21 ± 1 °C. b) Extrapolated relaxation times at zero piston displacement (y-intercepts of the fits in (a)), representing the intrinsic compressibility of each syringe material independent of the test fluid. All tests were conducted with n = 1.

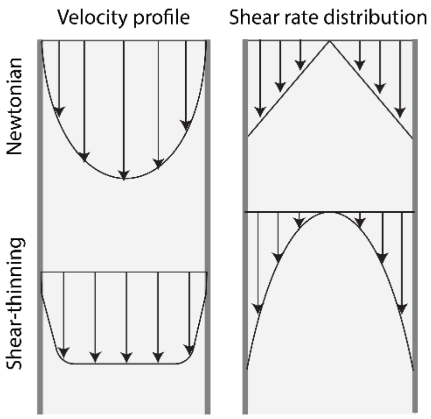

**Figure S13.** Velocity and shear rate profiles of a Newtonian (with parabolic velocity profile) and a shear-thinning materials (with plug velocity profile).

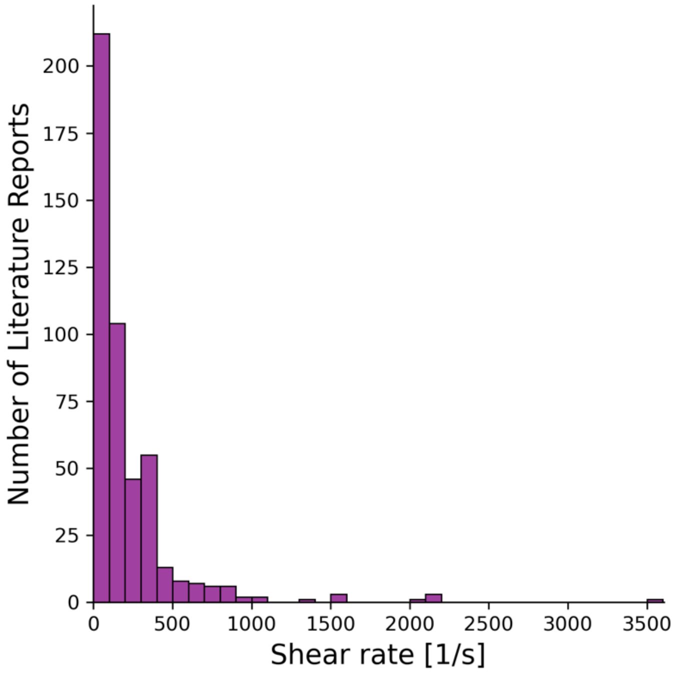

**Figure S14.** Shear-rate distribution extracted from 3D bioprinting literature based on reported bioprinting conditions.

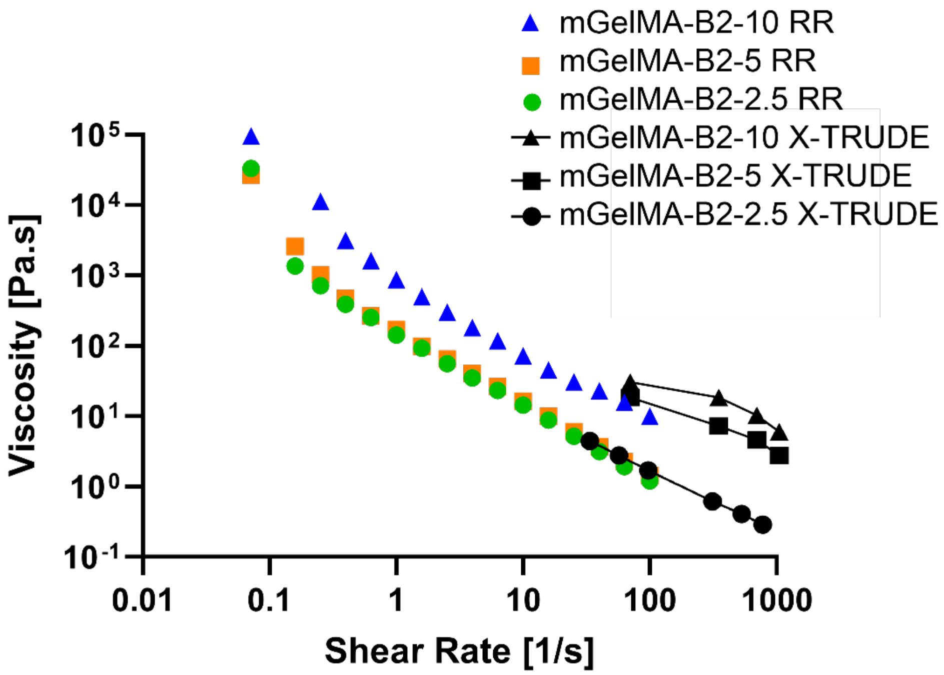

**Figure S15.** Flow curves acquired using RR and X-TRUDE for mGelMA formulations with second mGelMA batch (mGelMA-B2). All tests were conducted with n=2.

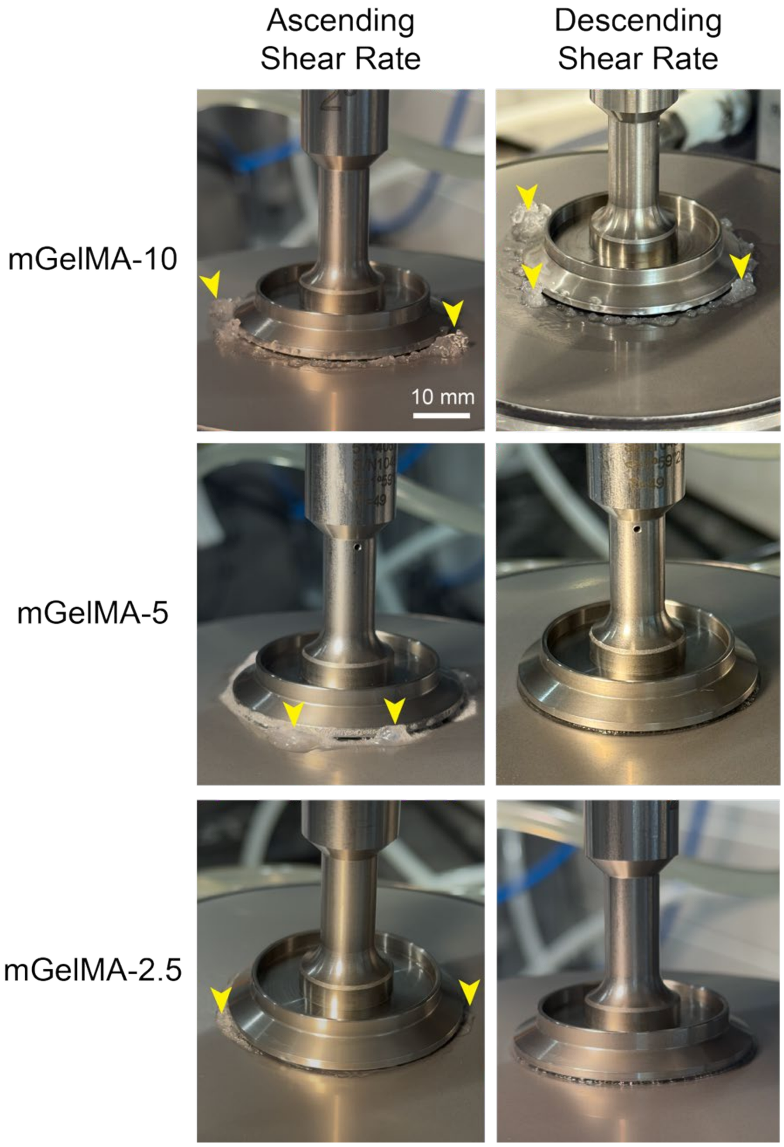

**Figure S16.** Expulsion of monolithic GelMA compositions made with second batch (mGelMA-B2) from the geometry-plate gap (indicated with yellow arrows) during flow sweep characterization with using rotational rheometry (RR). All mGelMA compositions were expelled during ascending shear rate implementation but only mGelMA-10 (both batch 1 and batch 2) was expelled during descending shear rate implementation.

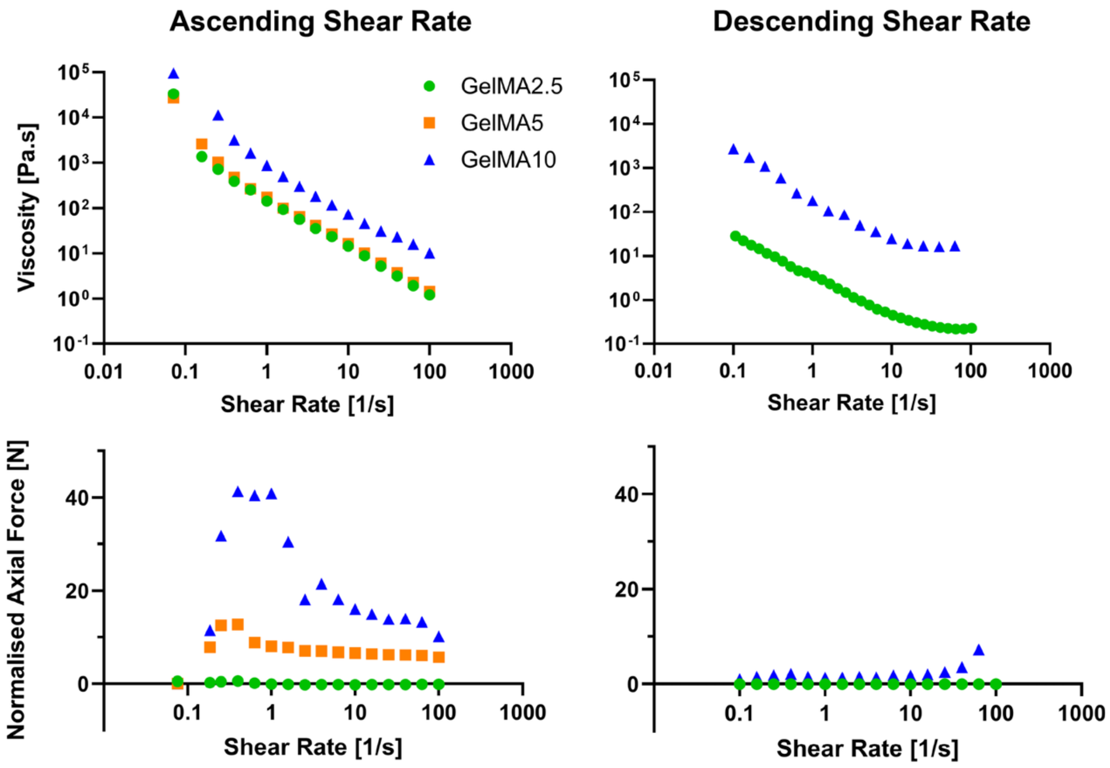

**Figure S17.** Flow curves for all mGelMA formulations acquired with RR using ascending and descending shear rate application, and their corresponding axial force response. All tests were conducted with n=1.

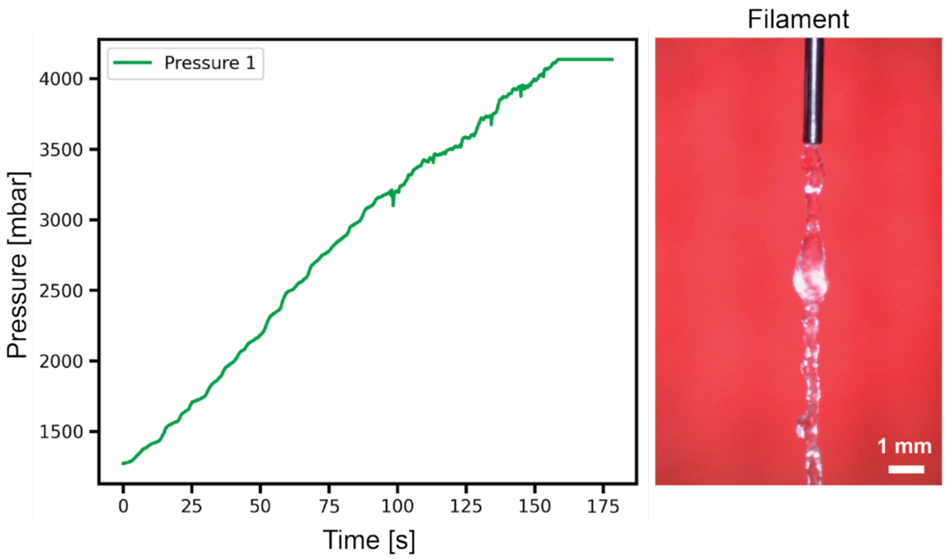

**Figure S18.** Pressure profile and corresponding filament of mGelMA-10 extruded through 25G cylindrical nozzle and 10 mms$^{-1}$ linear filament extrusion speed.

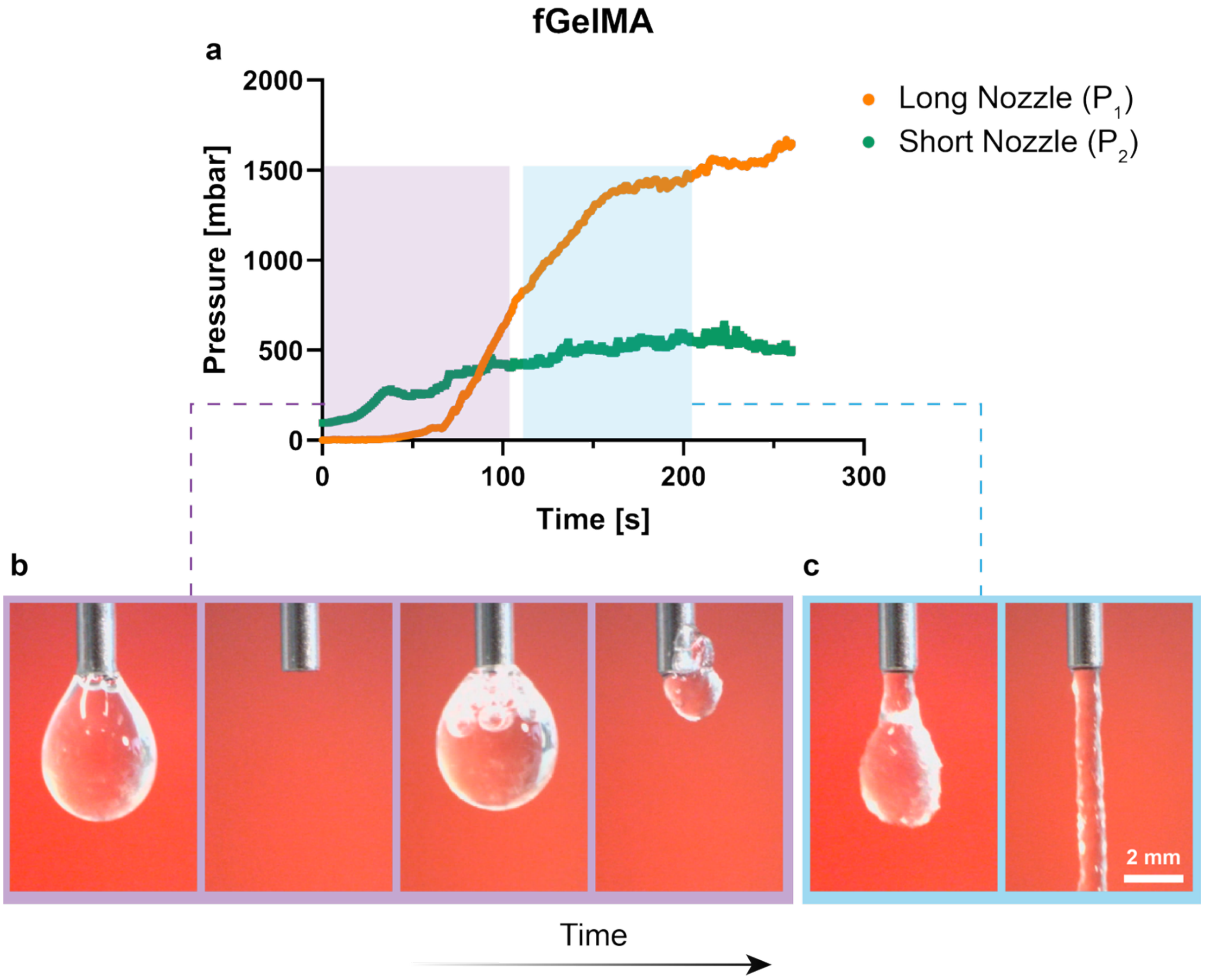

**Figure S19.** Representative example of the filter-pressing phenomenon observed during the first 60 seconds of gGelMA extrusion at 84.8 µL min$^{-1}$ and 21 ± 1 °C. a) Pressure curves obtained from syringe 1 equipped with a long nozzle (38.1 mm), and syringe 2 equipped with a short nozzle (~1 mm) during filter-pressing event. Corresponding extrudate photographs captured over time for the b) initial phase (purple region), and c) the later phase (blue region) of filter-pressing phenomenon.

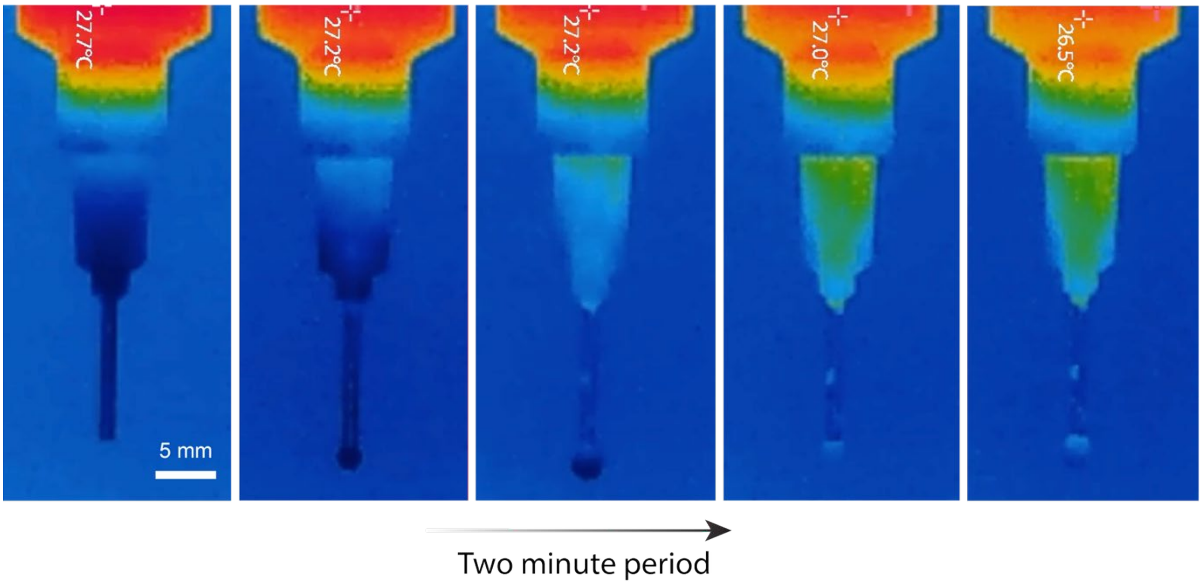

**Figure S20.** Thermal evolution of mGelMA-2.5 over two-minute period captured by IR thermal camera (mGelMA-2.5 preconditioned at 30 ± 1 °C extruded at chamber temperature of 21 ± 1 °C).

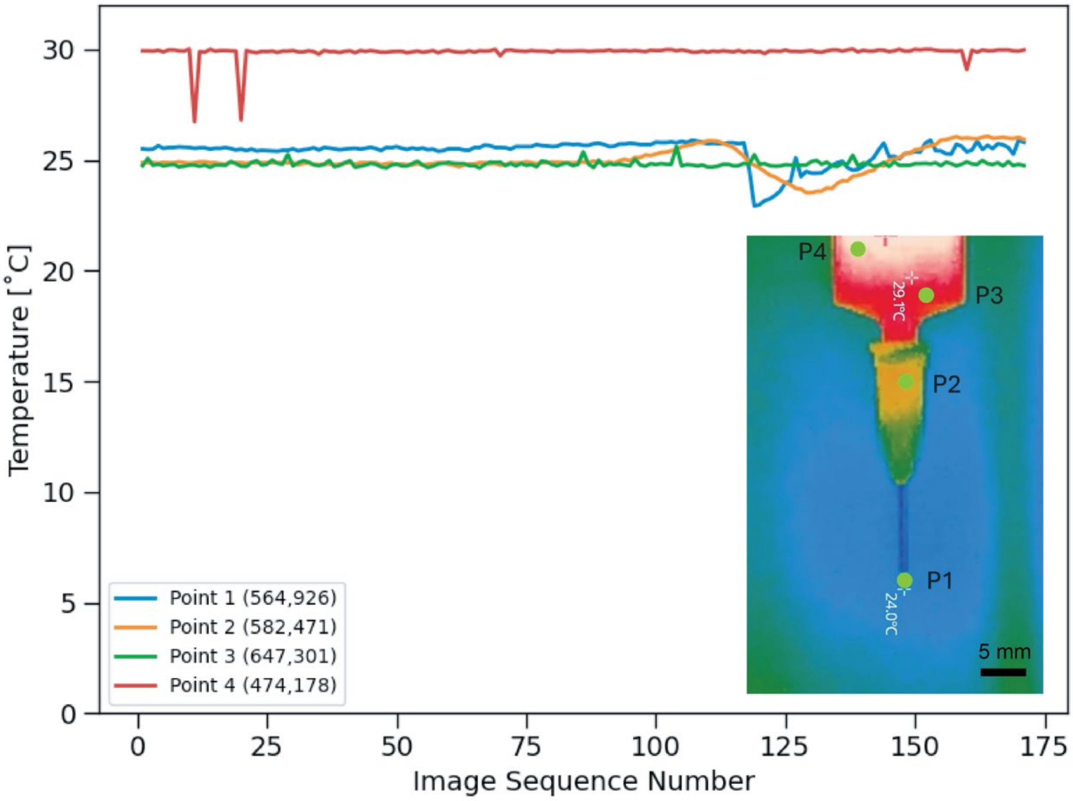

**Figure S21.** Thermal evolution of the four selected points over the two-minute extrusion course. Points of interest for temperature tracking are shown as P1, P2, P3, and P4.

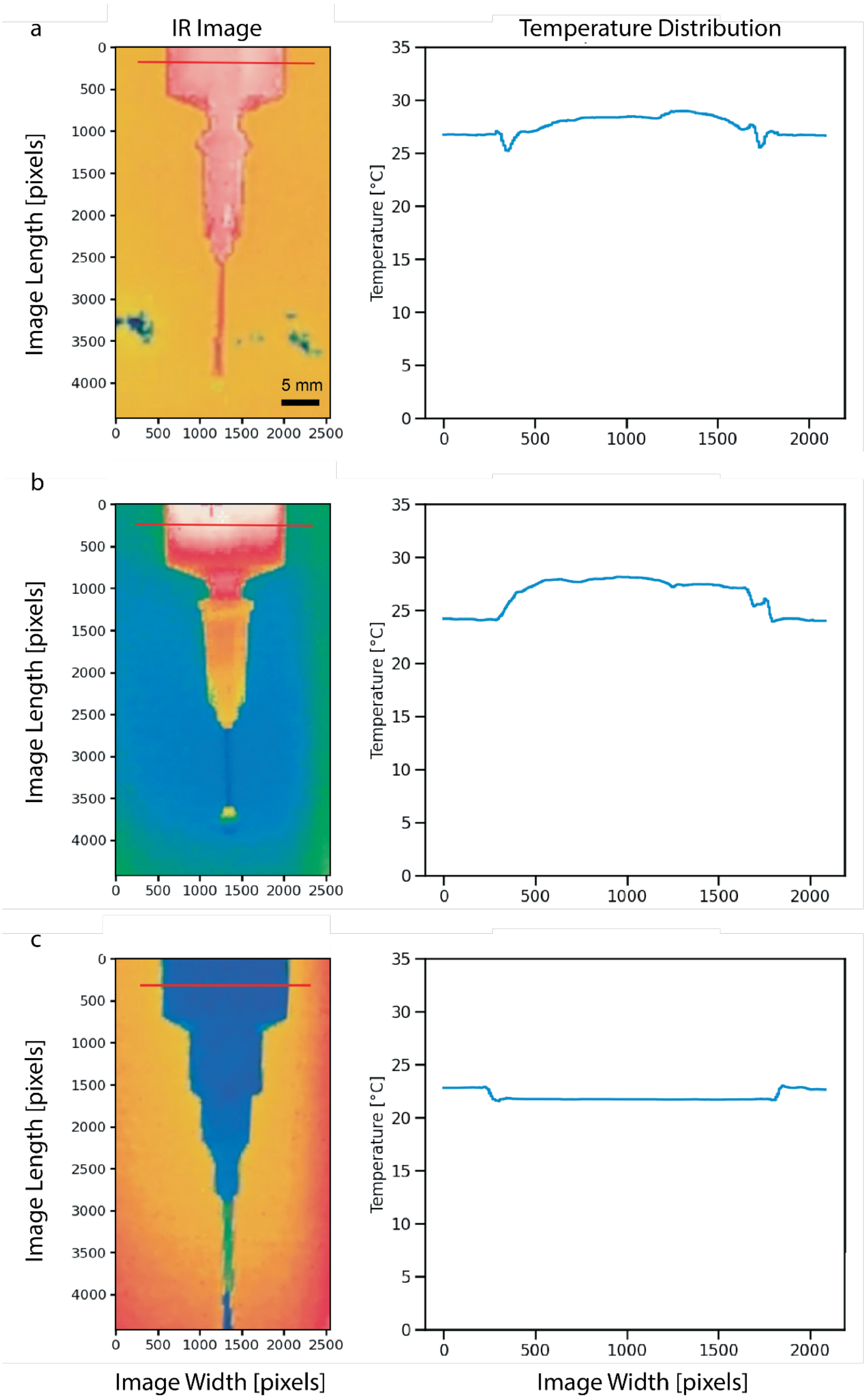
a
IR Image
Temperature Distribution
Image Length [pixels]
Temperature [°C]
5 mm
b
c
Image Width [pixels]
Image Width [pixels]

**Figure S22.** Infrared (IR) thermal images of syringe and extruded filament during mGelMA-2.5 extrusion (left) and corresponding temperature distribution (right) along the red reference line across the syringe width. a) Both the chamber and the material were maintained at 30 °C; b) The material was preconditioned to 30 °C, while chamber temperature was maintained at 21 ± 1 °C; c) both the chamber and the material were maintained at 21 ± 1 °C.

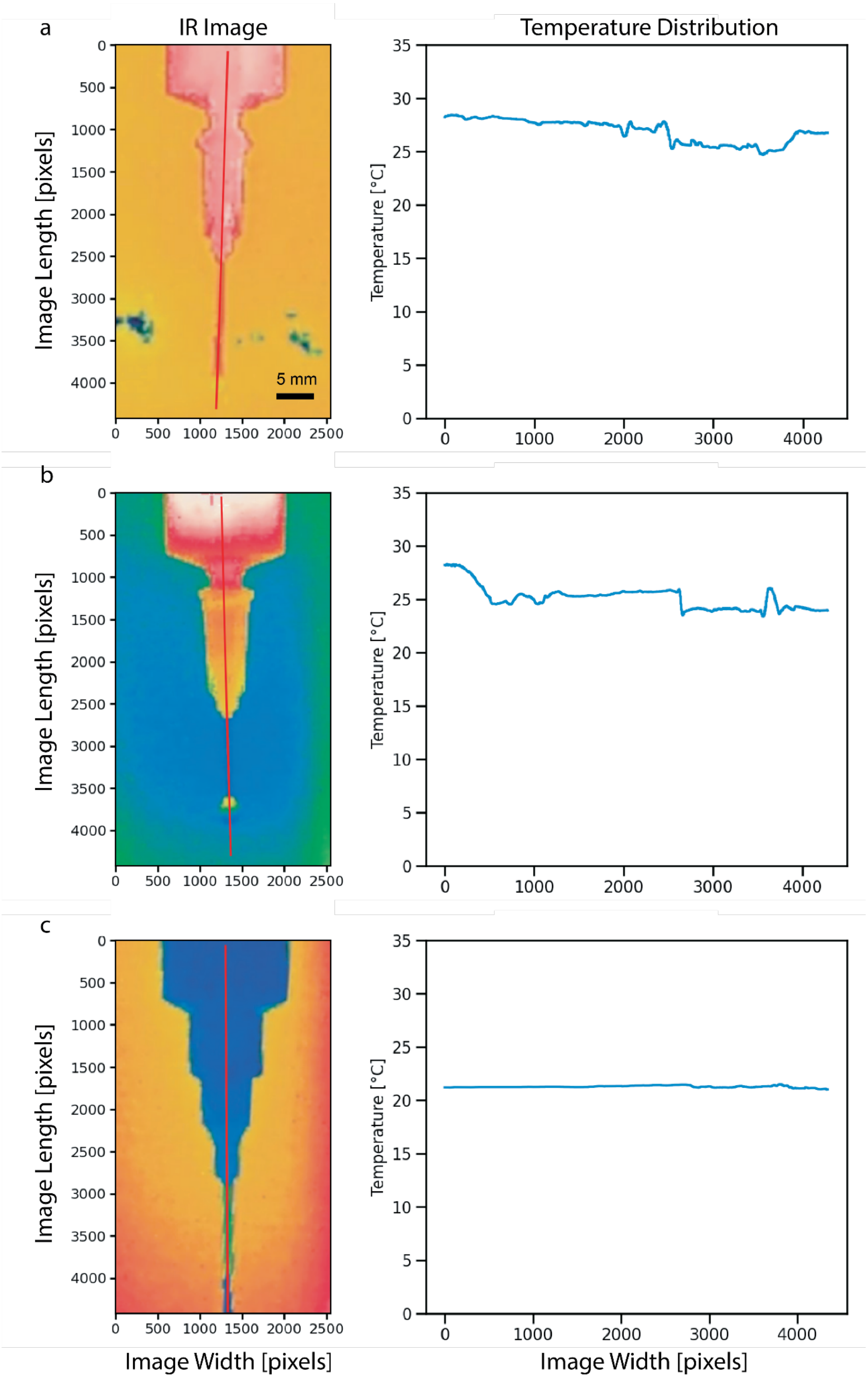
a
IR Image
Temperature Distribution
Image Length [pixels]
Temperature [°C]
5 mm
b
c
Image Width [pixels]
Image Width [pixels]

**Figure S23.** Infrared (IR) thermal images of syringe and extruded filament during mGelMA-2.5 extrusion (left) and corresponding temperature distribution (right) along the red reference line across the syringe and nozzle length. a) Both the chamber and the material were maintained at 30 °C; b) The material was preconditioned to 30 °C, while chamber temperature was maintained at 21 ± 1 °C; c) both the chamber and the material were maintained at 21 ± 1 °C.

#### 8. Supplementary Tables

**Table S1.** List of extrusion features obtained using X-TRUDE.

| Feature name | Identification | Material |
|---|---|---|
| Material exit | Initial Temporary Plateau | mGelMA |
| Filter-pressing | Low Long Needle Pressure vs Short Needle | gGelMA |
| Nozzle clogging | Pressure Spike | All |
| Filament morphology | Local Pressure Fluctuation | All |
| Shear thinning | Shear thinning index | All |
| Stress relaxation | Rheological calculations | All |

**Table S2.** Flow rates and the capillary sizes used for the calibration of the flow rates produced by X-TRUDE.

| Group name | Flow rate [$\mu$L.min$^{-1}$] | Capillary gauge |
|---|---|---|
| Low pressure | 10, 20, 50, 100, 200, 400, 600, 1000, 2000 | G25 |
| High pressure | 10, 20, 50, 100, 200, 400, 600, 1000, 2000 | G18 |
| High pressure | 10, 20, 50, 100, 200, 400, 600, 1000 | G22 |
| High pressure | 10, 20, 50, 100, 200 | G25 |

**Table S3.** Flow rates and the capillary sizes used for the calibration of viscosity measurements by X-TRUDE.

| Temperature | Flow rate [$\mu$L·min$^{-1}$] | Capillary gauge |
|---|---|---|
| 40 °C | 10, 20, 50, 100, 200, 400, 600, 1000, 2000 | G18 |
| 40 °C | 10, 20, 50, 100, 200, 400, 600, 1000 | G22 |
| 40 °C | 10, 20, 50, 100, 200 | G25 |
